# Ketamine reverses chronic corticosterone-induced changes in dendritic spine morphology in hippocampal but not cortical neurons in vitro

**DOI:** 10.64898/2026.11.27.769436

**Authors:** Michał Wilczkowski, Krzysztof Kączek, Piotr Chmielarz, Patrycja Mieszczak, Klaudiusz Górka, Anna Alwani, Agnieszka Zelek-Molik, Irena Nalepa

## Abstract

Chronically elevated glucocorticoids, such as corticosterone (CORT), reduce dendritic spine density in the prefrontal cortex and hippocampus, and the rapid-acting antidepressant ketamine (KET) reverses this spine loss in the prefrontal cortex. Less is known about how CORT affects spine morphology, whether these effects differ between the cortex and hippocampus, and whether KET counteracts them. Most studies used stress models or systemic CORT, making it difficult to distinguish direct and indirect effects of CORT on neurons. We therefore used primary mouse cortical and hippocampal cultures, which isolate these direct effects. We found that chronic CORT (0.25 µM, 72 h) altered spine morphology differently in each culture. In cortical neurons, CORT narrowed spine heads and increased the length-to-head-width ratio, and the selective glucocorticoid receptor (GR) modulator CORT108297 abolished the CORT-induced narrowing of spine heads. By contrast, in hippocampal neurons, CORT reduced spine length, head width, and spine area, making dendritic spines proportionally smaller. Because dendritic spine structure and network activity shape each other, we also recorded spontaneous Ca²⁺ activity. The regional divergence observed in spine structure also extended to network activity: CORT shortened the intervals between Ca²⁺ events in hippocampal but not in cortical cultures. KET (10 µM, 24 h) did not counteract CORT effects in cortical cultures, whereas in hippocampal cultures it reversed the CORT-induced reduction in spine size. In both cultures, KET reduced network synchronization independently of CORT. Thus, CORT remodels dendritic spines in a region-specific manner, and KET counteracts its effects on spine morphology only in hippocampal neurons.

## Introduction

A remarkable feature of the brain is its ability to reorganize. The basic cellular components that underlie brain plasticity are dendritic spines – membranous protrusions that constitute the postsynaptic site of most excitatory connections [1, 2]. Dendritic spines are highly motile; they can change shape rapidly and may also be formed and eliminated [3]. Crucially, their structure reflects synaptic strength, as spine head volume positively correlates with α-amino-3-hydroxy-5-methyl-4-isoxazolepropionic acid receptor (AMPAR) abundance and synapse stability [4, 5]. Structural remodeling of spines therefore influences the strength and connectivity of neural circuits, and its dysregulation has been linked to neuropsychiatric disorders [6]. Characterizing how specific factors reshape spine morphology could thus help to understand how circuit function becomes dysregulated. Chronic stress, acting largely through its principal mediator corticosterone (CORT), is one such factor [7].

CORT acts through the high-affinity mineralocorticoid receptor (MR) and the lower-affinity GR, which is recruited at elevated CORT levels [8]. When its physiological oscillations are undisturbed, CORT promotes synaptic plasticity [9, 10]. Circadian CORT peaks drive learning-dependent spine formation, while troughs consolidate the subset of those newly formed spines that supports long-term memory [9]. However, when chronically in excess, CORT drives excessive dendritic spine elimination, including the loss of learning-associated and long-lasting spines, in vulnerable regions (primarily the prefrontal cortex and hippocampus), accompanied by cognitive impairment and depressive-like behaviors [9, 11, 12]. Beyond spine elimination, chronically elevated CORT has been shown to weaken synaptic connections, shifting spines of primary hippocampal neurons toward immature, small-headed phenotype; however, such in vitro data remain scarce and limited to the hippocampus [13]. Most evidence about CORT effects on dendritic spines comes instead from multifactorial stress models or systemic CORT [9, 11, 14]. In these models, CORT can alter cortical and hippocampal neurons indirectly, for example through altered input from other brain regions, such as the thalamus, or through peripheral changes; hence, the specific, local contribution of CORT itself is difficult to disentangle [15]. In addition, most studies have addressed spine formation and elimination rather than morphology, and the few that examined morphology relied mainly on classification into spine subtypes or on single size measures [11, 16]. It therefore remains unclear whether CORT reshapes spine morphology, and thus synaptic strength, differently across vulnerable regions, and whether these effects involve GR.

Disentangling the effects of CORT on spine morphology is critical, as disturbed spine structure may contribute to the pathophysiology of depression [17]. Postmortem and in vivo imaging studies revealed reduced synapse density in the prefrontal cortex of depressed patients, and cortisol, the human counterpart of rodent CORT, is dysregulated in a subset of them [18–20]. Restoring synaptic connectivity is therefore an important therapeutic goal, all the more so because conventional antidepressants act with a delay of weeks and are ineffective in a large proportion of depressed individuals [17, 21]. Ketamine (KET), an N-methyl-D-aspartate receptor (NMDAR) antagonist, however, largely overcomes both limitations: a single dose produces rapid and sustained antidepressant effects, including in treatment-resistant patients [22, 23]. A study connecting KET, CORT and dendritic spines showed that KET restored spines eliminated by chronic CORT and promoted de novo spine formation at new sites in the prefrontal cortex. Notably, although KET restored neuronal ensemble activity within hours, before spine formation, optical deletion of the newly formed spines abolished its long-term effects on prefrontal activity [12]. However, data on how KET regulates spine morphology after chronic CORT exposure are lacking. Whether KET similarly restores CORT-disrupted spines in other vulnerable structures, such as the hippocampus, also remains unknown.

To address these gaps, we used primary mouse cortical and hippocampal cultures, which allow the local effects of CORT to be isolated from these indirect effects. Because the standard B27 supplement contains CORT, cultures were transferred to a CORT-free medium before treatment, so that CORT exposure was fully controlled. First, we selected a CORT concentration that engaged glucocorticoid signaling, as indicated by increased expression of the GR target gene *Fkbp5*, without reducing cell viability or total dendritic length. Second, we characterized how chronic CORT affects spine density, spine morphology (length, head width, area, and length-to-head-width ratio), and PSD-95 puncta in cortical and hippocampal neurons, and whether these effects depend on GR, using the selective GR modulator CORT108297 (C108). Third, we examined whether KET, applied during the last 24 h of CORT treatment, counteracts CORT-induced changes in spine morphology in both cultures. Finally, because spine morphology and network activity are bidirectionally linked, with spine structure shaping circuit connectivity and information processing and network activity in turn remodeling spines, we used Ca²⁺ imaging to test whether the effects of CORT and KET on spines were accompanied by changes in spontaneous network activity and synchronization.

## Materials and methods

### Primary cell culture

Primary cortical and hippocampal cultures were prepared from E16–17 Albino Swiss mouse embryos (sex was not determined). Pregnant dams were euthanized by cervical dislocation, and the embryos were removed surgically. The embryos were placed in a plastic dish filled with ice-cold, Ca²⁺/Mg²⁺-free Hanks’ Balanced Salt Solution (HBSS; Thermo Fisher Scientific, cat. no. 14175095). The cortices and hippocampi were dissected from each brain and transferred to separate tubes containing ice-cold HBSS, with tissue from all embryos pooled per structure. The HBSS was then replaced with trypsin solution (2.5 mg/ml; MP Biomedicals, cat. no. TRP0004.25), and the tissue was digested for 15 min at 37 °C. After digestion, the trypsin solution was removed, and the tissue was washed twice with HBSS containing 10% (v/v) horse serum (Thermo Fisher Scientific, cat. no. 16050122) and 0.006% (w/v) DNase I (Roche, cat. no. 11284932001). The tissue was then triturated in fresh HBSS with horse serum and DNase I using a 1-ml pipette tip until a homogeneous suspension was obtained. The suspensions were transferred to 15-ml conical tubes (one per structure) and centrifuged at 200 × g for 3 min. The cell pellets were resuspended in plating medium consisting of Neurobasal medium (Thermo Fisher Scientific, cat. no. 12348017) supplemented with 2% (v/v) B27 (Thermo Fisher Scientific, cat. no. 17504044) and 0.25% (v/v) GlutaMAX (Thermo Fisher Scientific, cat. no. 35050061). Cells were counted with an automated cell counter (TC20; Bio-Rad) and plated on poly-D-lysine-coated coverslips or plates (Supplementary Methods) at 1.2 × 10⁵ cells per coverslip, 4 × 10⁴ cells per well of a 96-well plate, and 1 × 10⁶ cells per well of a 6-well plate. Cultures were maintained at 37 °C in a humidified atmosphere with 5% CO₂.

### Medium changes

Primary cortical and hippocampal cultures were plated and maintained for the first 5/6 days in vitro (DIV) in Neurobasal medium supplemented with B27 and GlutaMAX (see *Primary cell culture*). However, B27 contains CORT, the independent variable in this study. To control the CORT concentration in the culture medium, the medium was therefore changed four times between plating and the start of the experiments (i.e., the onset of CORT treatment). The first three changes replaced one-third of the medium, whereas the fourth, always performed one day before the experiment, replaced three-quarters of the medium. The replacement medium consisted of Minimum Essential Medium (MEM; Thermo Fisher Scientific, cat. no. 51200038) supplemented with 0.45% (w/v) D-glucose (added from a 45% solution; Sigma-Aldrich, cat. no. G8769), 1% (v/v) N2 supplement (Thermo Fisher Scientific, cat. no. 17502048), 0.25% (v/v) GlutaMAX (Thermo Fisher Scientific, cat. no. 35050061), 0.22 mM sodium pyruvate (Sigma-Aldrich, cat. no. P2256), and 10 mM HEPES (Thermo Fisher Scientific, cat. no. 15630080). In the experiment comparing the effects of culture media on spine morphology parameters, the medium was changed four times. In the MEM/N2 groups, medium changes were performed as described above, whereas in the Neurobasal/B27 groups, the medium was changed on the same schedule but with fresh Neurobasal/B27. CORT (0.25 µM) was added at DIV 18 or 19.

### Drug treatments

The following drugs were used: corticosterone (CORT; Sigma-Aldrich, cat. no. 27840), ketamine hydrochloride (KET; Tocris, cat. no. 3131), and the selective GR modulator CORT108297 (C108; MedChemExpress, cat. no. HY-15710). CORT and C108 were dissolved in dimethyl sulfoxide (DMSO; Sigma-Aldrich, cat. no. 276855), and KET was dissolved in MEM. CORT was used at 0.25, 0.5, and 1 µM. CORT stock solutions were prepared in DMSO at 10,000× the final concentrations and diluted in two 100-fold steps: first in MEM, then directly in the culture medium. The final DMSO concentration was therefore 0.01% (v/v). Vehicle-treated controls received DMSO diluted in the same manner. KET was used at 10 µM, diluted 100-fold in the culture medium from a 1 mM stock in MEM; groups not receiving KET received an equal volume of MEM. C108 was used at 100 nM, prepared from a 1 mM DMSO stock in the same two-step manner as CORT. In experiments combining CORT and C108, all groups received 0.02% DMSO. Cells were treated with CORT for 72 h; KET was added for the last 24 h of CORT treatment, and C108 was applied together with CORT. CORT treatment started at DIV 18 or 19. Initial concentrations of CORT and KET were selected based on previous studies [13, 24, 25]. C108 was used at 100 nM, approximately 100-fold above its Ki for GR (0.9 nM) and approximately 100-fold below the Ki values reported for MR [26, 27]. The final CORT concentration was chosen based on dose–response experiments. Subsequent experiments used 0.25 µM CORT only, as it was the lowest tested concentration that increased *Fkbp5* expression and altered spine morphology without reducing cell viability or total dendritic length (Figs. 1 and 2). The 72 h incubation with CORT was likewise based on previous work in primary cortical neurons, in which 3 days (72 h) of CORT exposure was sufficient to reduce mitochondrial GR and Bcl-2 levels; a comparable decrease in mitochondrial Bcl-2 was observed in the prefrontal cortex of rats after 21 days of CORT administration in the drinking water [28].

**Fig. 1.**
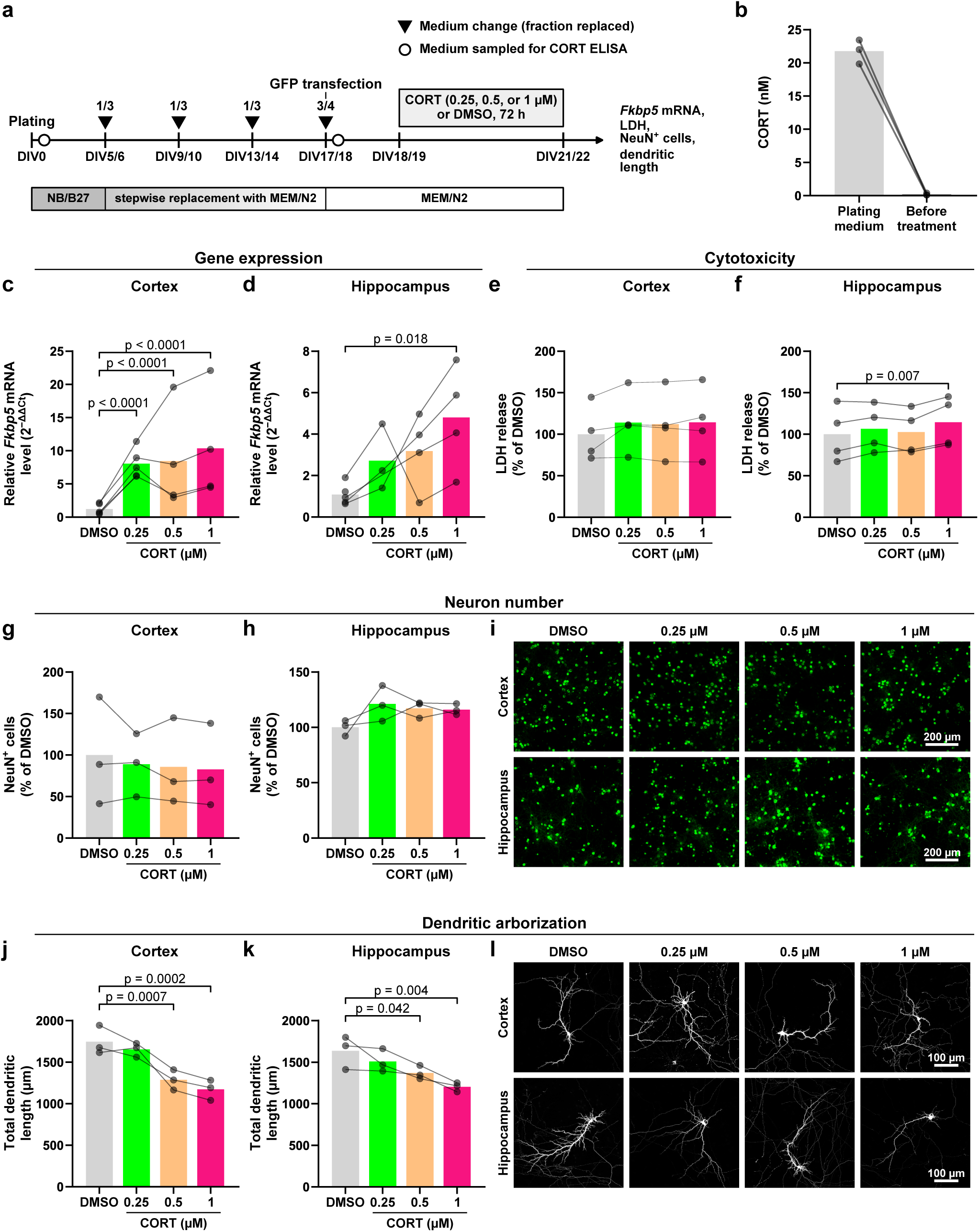
Validation of the culture model and choice of CORT concentration. **a** Experimental timeline. Cultures were plated in Neurobasal/B27 (NB/B27), which was replaced stepwise with CORT-free MEM/N2 by four partial medium changes (fraction of medium replaced is indicated above the arrowheads). Neurons were transfected with GFP at DIV17/18, and CORT (0.25, 0.5, or 1 µM) or vehicle (DMSO) was applied for 72 h from DIV18/19. Open circles mark medium sampling for the CORT ELISA. **b** CORT concentration in the plating medium and in the medium collected after the fourth medium change, one day before treatment (n = 3 cultures). **c, d** *Fkbp5* mRNA levels in cortical (**c**) and hippocampal (**d**) cultures relative to DMSO (n = 5, 5, 4, and 4 cultures in **c** and 5, 3, 4, and 4 cultures in **d** for DMSO, 0.25, 0.5, and 1 µM, respectively). In **c** and **d**, bars represent arithmetic means of 2^−ΔΔCt^ values, whereas statistical analyses were performed on log-transformed values and geometric mean fold changes are given in the text. **e, f** LDH release from cortical (**e**) and hippocampal (**f**) cultures (n = 4 cultures). **g, h** Number of NeuN⁺ cells in cortical (**g**) and hippocampal (**h**) cultures (one well per condition, 23 fields per well; n = 3 cultures). **i** Representative images of NeuN immunostaining. Fields were selected so that the number of NeuN⁺ cells relative to DMSO matched the group means. **j, k** Total dendritic length of GFP-transfected cortical (**j**) and hippocampal (**k**) neurons. In total, 49, 47, 50, and 53 cortical neurons and 34, 40, 38, and 32 hippocampal neurons were analyzed for DMSO, 0.25, 0.5, and 1 µM CORT, respectively; n = 3 cultures. **l** Representative maximum-intensity projections of GFP-transfected neurons. Bars represent means, points represent individual cultures, and lines connect values from the same culture. p values from Dunnett’s post hoc test vs DMSO are shown for significant comparisons. Scale bars: 200 µm (**i**), 100 µm (**l**)

**Fig. 2.**
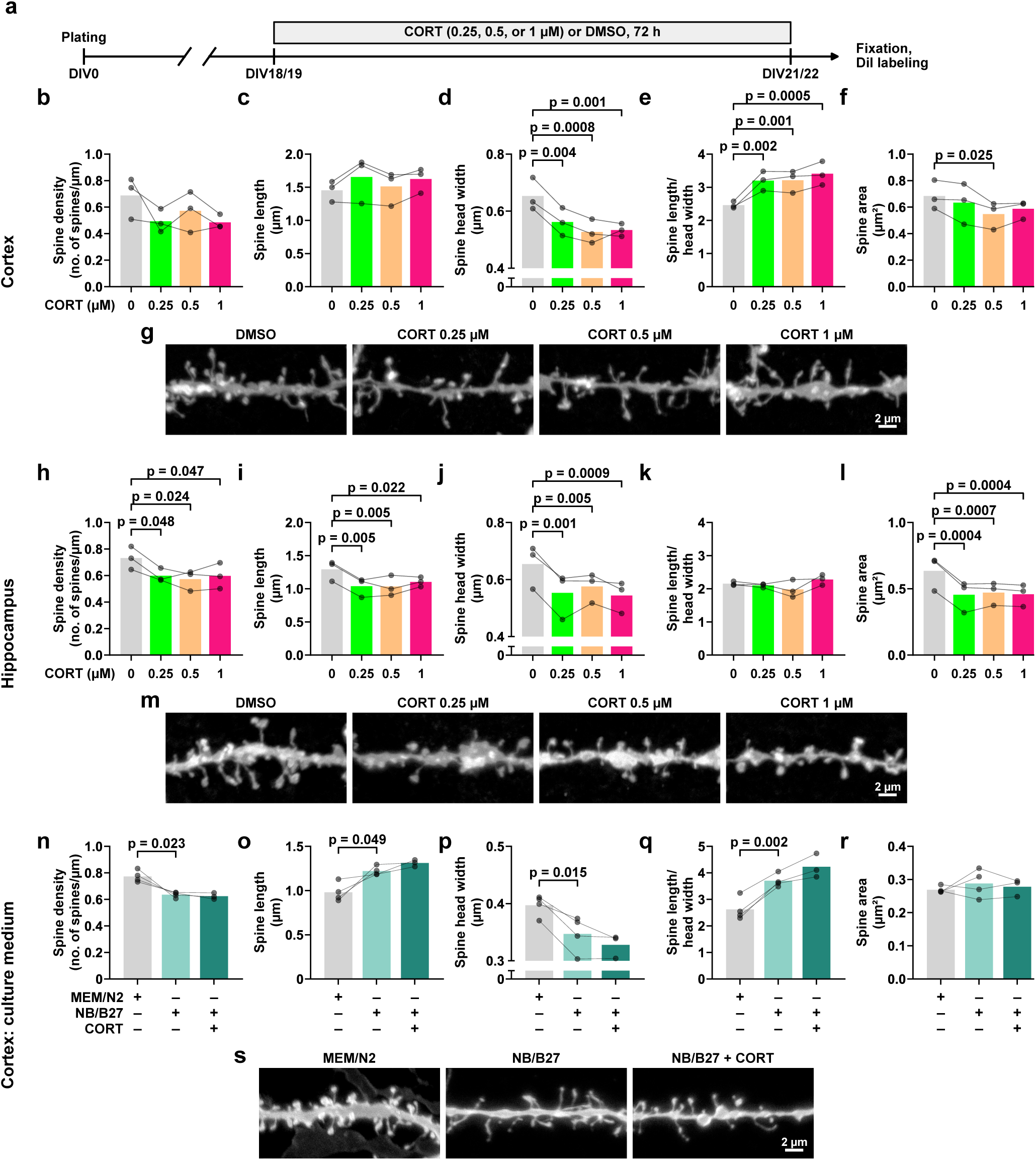
Chronic CORT alters dendritic spine morphology in a region-specific manner, and culture medium affects baseline spine morphology. **a** Experimental timeline. Cultures were treated with CORT (0.25, 0.5, or 1 µM) or DMSO for 72 h from DIV18/19 and fixed at DIV21/22. **b–f** Spine density (**b**), spine length (**c**), spine head width (**d**), length-to-head-width ratio (**e**), and spine area (**f**) in cortical neurons. **g** Representative DiI-labeled dendritic segments (20 µm) of cortical neurons. **h–l** The same parameters in hippocampal neurons. **m** Representative dendritic segments of hippocampal neurons. In total, 5247 spines on 9292 µm of dendrites were analyzed in cortical cultures and 6538 spines on 10,610 µm of dendrites in hippocampal cultures; n = 3 cultures. p values from Dunnett’s post hoc test vs DMSO are shown for significant comparisons. **n–r** The same parameters in cortical neurons maintained in MEM/N2 (standard protocol of this study) or in Neurobasal/B27 (NB/B27) without or with 0.25 µM CORT. In total, 4978 spines on 7437 µm of dendrites were analyzed; n = 4 cultures (MEM/N2 and NB/B27) and 3 cultures (NB/B27 + CORT). p values from paired t-tests (MEM/N2 vs NB/B27) are shown for significant comparisons. **s** Representative dendritic segments for **n–r**. In **g**, **m**, and **s**, segments were taken from one neuron per condition and chosen from straight dendritic stretches without overlapping neurites. Bars represent means, points represent individual cultures, and lines connect values from the same culture. Scale bars: 2 µm

### Cell transfection

Primary neurons were transfected at DIV 17–18 with the pCDH-hSYN-MCS-T2A-GFP plasmid (System Biosciences) by magnetofection, using NeuroMag (OZ Biosciences, cat. no. KC30800) according to the manufacturer’s protocol. First, plasmid DNA (0.5 µg per well of a 96-well plate), diluted in MEM, was mixed with NeuroMag (1 µl per well) by pipetting and incubated for 20 min at room temperature. The DNA–NeuroMag complexes were then added to the culture medium in a volume of 50 µl per well, and the culture plate was placed on a magnetic plate in a humidified incubator (37 °C, 5% CO₂) for 18 min. Finally, the fourth medium change was performed (see section *Medium changes*).

### Fixation and DiI staining

Primary cortical and hippocampal cultures were fixed at DIV 18 (cell culture characterization), 21, or 22, depending on the experiment, with pre-warmed (37 °C) paraformaldehyde (PFA) in PBS. After removal of the culture medium, cells were incubated for 15 min at room temperature with 1.5% PFA for spine morphology and postsynaptic density protein 95 (PSD-95) experiments or 4% PFA for all other immunofluorescence experiments. The PFA was then removed, and the cells were washed once with PBS and stored in fresh PBS at 4 °C until further processing.

Dendritic spines of cortical and hippocampal neurons were visualized with the lipophilic dye 1,1′-dioctadecyl-3,3,3′,3′-tetramethylindocarbocyanine perchlorate (DiI; Sigma-Aldrich, cat. no. 42364). Depending on the experiment, DiI staining was performed alone or after immunofluorescence staining. DiI crystals were suspended in PBS (20 µg/ml) and sonicated for 1–2 h until a homogeneous suspension was obtained. The suspension was then diluted 10-fold in fresh PBS and added at 1.2 ml per well of a 12-well plate (coverslips) or 120 µl per well of a 96-well plate. Cells were incubated with DiI for 20 min at 37 °C, after which the DiI solution was removed and the cells were washed three times with PBS. The 96-well plates were kept at room temperature overnight to allow dye diffusion and then stored at 4 °C until further processing. Coverslips were mounted on microscope slides with ProLong Gold Antifade Mountant (Thermo Fisher Scientific, cat. no. P36930), dried for 1 h at room temperature, and stored at −20 °C until imaging.

### Imaging and analysis of dendritic spines

Images of DiI-stained dendritic spines were acquired with a Leica TCS SP8 X confocal microscope (Leica Microsystems) using an HC PL APO CS2 100×/1.40 oil-immersion objective. DiI was excited at 555–557 nm, Z-stacks were acquired at 1024 × 1024 pixels with a 1.63× zoom (pixel size, 0.07 µm, satisfying the Nyquist criterion), a z-step of 0.40 µm, and 4× line averaging. Acquisition settings were kept constant across conditions within each biological replicate and were adjusted between replicates to account for differences in labeling intensity. DiI labeled a sparse, random subset of neurons; at least eight labeled neurons per well were selected at random and imaged. Spines on secondary and tertiary dendrites were analyzed to avoid differences in spine structure related to dendritic branch order. Spines were analyzed semi-automatically in maximum-intensity projections using SpineMagick (international patent no. WO/2013/021001) to quantify spine density (number of spines per 1 µm of dendrite), spine length, head width, length-to-head-width ratio, and spine area [29]. Analyses were performed by experimenters blinded to treatment. Absolute values of spine parameters differ between the concentration–response experiment, performed on coverslips mounted in ProLong Gold, and the remaining experiments, performed in 96-well imaging plates, likely reflecting differences in growth substrate, mounting medium, and imaging conditions. Each experiment was, however, analyzed separately in a randomized block design, with all conditions processed in parallel within each culture, and data from different experiments were not pooled. The number of dendrites and spines analyzed per experiment is given in the figure legends. Representative neurons shown in figures were selected to approximate the group means of the analyzed spine parameters, and the brightness and contrast of the displayed dendritic segments were adjusted linearly and independently for each image for display purposes only.

### Immunofluorescence staining

After fixation (see *Fixation and DiI staining*), cells were permeabilized with 0.3% (v/v) Tween-20 in PBS for 15 min and then blocked with 10% (v/v) normal horse serum in PBS containing 0.3% Tween-20 for 1 h at room temperature. After one wash with PBS, cells were incubated overnight at 4 °C with the following primary antibodies diluted in PBS containing 1% (v/v) normal horse serum and 0.3% Tween-20: mouse anti-PSD-95 (Abcam, cat. no. ab2723; 1:250), mouse anti-NeuN (Abcam, cat. no. ab104224; 1:1000), rabbit anti-S100B (ABclonal, cat. no. A19108; 1:500), chicken anti-MAP2 (Abcam, cat. no. ab5392; 1:5000), and goat anti-IBA1 (Abcam, cat. no. ab5076; 1:1000). After three washes with PBS, cells were incubated for 2 h at room temperature with cross-adsorbed, Alexa Fluor-conjugated secondary antibodies: donkey anti-mouse Alexa Fluor 488 (Thermo Fisher Scientific, cat. no. A-21202; 1:250) and 647 (Thermo Fisher Scientific, cat. no. A-31571; 1:250), donkey anti-rabbit Alexa Fluor 488 (Thermo Fisher Scientific, cat. no. A-32790TR; 1:250) and 647 (Thermo Fisher Scientific, cat. no. A-31573; 1:250), donkey anti-chicken Alexa Fluor 568 (Thermo Fisher Scientific, cat. no. A78950; 1:250), and donkey anti-goat Alexa Fluor 647 (Thermo Fisher Scientific, cat. no. A-21447; 1:250). Cells were then washed three times with PBS, incubated with Hoechst 33342 (Thermo Fisher Scientific, cat. no. H3570; 1:20,000) for 20 min at room temperature, washed once with PBS, and stored in fresh PBS at 4 °C until further processing.

### Imaging and analysis of dendritic arborization

Images of GFP-transfected neurons were acquired with a Leica TCS SP8 X confocal microscope (Leica Microsystems) using an HC PL APO CS2 20×/0.75 dry objective. GFP was excited at 488 nm, Z-stacks were acquired at 1024 × 1024 pixels (pixel size, 0.48 µm) with a z-step of 1 µm, 1.2× zoom and 8× line averaging. Laser power, detector gain, and all other acquisition settings were kept constant across conditions within each biological replicate and were adjusted between replicates to account for differences in labeling intensity. All GFP-positive neurons in each well were imaged. Maximum-intensity projections were analyzed semi-automatically in ImageJ (Fiji) using the SNT plugin [30] to quantify total dendritic length. The number of neurons analyzed per group is given in the figure legends.

### Imaging and analysis of PSD-95 puncta

Images of PSD-95 immunostaining in dendrites and dendritic spines were acquired with a Leica TCS SP8 X confocal microscope (Leica Microsystems) using an HC PL APO CS2 100×/1.40 oil-immersion objective. DiI, used to visualize dendrites and dendritic spines, was excited at 555 nm, and Alexa Fluor 647, used to detect PSD-95, was excited at 653 nm. Multichannel images were acquired sequentially to avoid spectral bleed-through. Z-stacks were acquired at 1024 × 1024 pixels (pixel size, 0.07 µm) with a z-step of 0.40 µm. Laser power, detector gain, and all other acquisition settings were set to avoid pixel saturation and were kept constant across each experiment.

PSD-95 puncta were quantified in ImageJ (Fiji) in selected secondary and tertiary dendrites of cortical and hippocampal neurons. Binary masks generated from DiI images were applied to the corresponding PSD-95 images to retain only the PSD-95 signal within DiI-labeled dendrites and spines, excluding the soma. Maximum-intensity projections of the masked images were thresholded using the Moments algorithm, applied to all images within a series. PSD-95 puncta in manually selected secondary and/or tertiary dendrites were analyzed using Analyze Particles, and dendrite length was measured with the Segmented Line tool. Puncta density (number of puncta per µm of dendrite), size, and mean fluorescence intensity were compared between DMSO-and CORT-treated cultures. Representative images were cropped for display without further intensity adjustments.

### Widefield imaging and cell counting

Images of NeuN-, MAP2-, S100B-, and Iba1-stained cells were acquired with an inverted widefield fluorescence microscope (Leica Thunder DMI8) using an HC PL FLUOTAR 10×/0.32 objective. Excitation wavelengths were 470 nm for NeuN, 550 nm for MAP2, 470 or 640 nm for S100B (depending on the experiment), 640 nm for Iba1 and 405 nm for Hoechst 33342. For multichannel imaging, secondary antibodies were combined to avoid spectral bleed-through. Representative images of MAP2, S100B, and Iba1 were acquired at 1024 × 1024 pixels. Images for NeuN^+^ cell counting were acquired at 2048 × 2048 pixels with 470 nm excitation and a 700 ms exposure time. Images used for cell counting were processed with Large Volume Computational Clearing (LVCC; Leica) using identical settings for all images, and the processed images were analyzed. Within each experiment, acquisition parameters were identical across conditions.

NeuN^+^ cells were counted in Fiji (ImageJ). To determine the number of NeuN ^+^ cells, images were thresholded using the MaxEntropy method, touching cells were separated using the watershed algorithm, and NeuN^+^ cells were counted in the binarized images using Analyze Particles, excluding objects touching the image border. The number of NeuN⁺ cells was determined in one well per condition (23 fields per well) in each culture. The proportions of NeuN⁺ and S100B⁺ cells were determined using the Trainable Weka Segmentation plugin [31] (Supplementary Methods). Representative NeuN images shown in the figure were selected so that the number of NeuN⁺ nuclei relative to DMSO matched the group means, and their brightness was adjusted linearly and independently for each image.

### Calcium imaging and analysis

Spontaneous Ca²⁺ activity was imaged using the synthetic Ca²⁺ indicator Cal-520 AM (AAT Bioquest, cat. no. 21131). Cal-520 AM was dissolved in DMSO to 2 mM, aliquoted, and stored at −20 °C. On the day of the experiment, the stock was diluted 5-fold in Ca²⁺/Mg²⁺-free HBSS and then 100-fold in culture medium, yielding a final concentration of 4 µM (0.2% DMSO). In accordance with the manufacturer’s protocol, cells were loaded for 75 min at 37 °C in a humidified atmosphere with 5% CO₂. Pluronic F-127, which is used to increase the aqueous solubility of Cal-520 AM, was omitted, as it can alter cytoplasmic Ca²⁺ regulation in neurons [32]. The loading medium was then replaced with conditioned medium collected from parallel, unloaded wells containing cells from the same culture and subjected to the same treatment, so that the treatment conditions were maintained during imaging.

Images of live, dye-loaded cells were acquired using an Opera Phenix high-content screening system (Revvity) with an environmental chamber set to 37 °C and 5% CO₂. Imaging was performed in non-confocal (widefield) epifluorescence mode with a 10× objective (NA 0.3) and an Andor Zyla sCMOS camera with 4 × 4 binning (540 × 540 pixels, 2.37 µm/pixel, field of view 1.28 × 1.28 mm). Cal-520 fluorescence was excited at 488 nm with an exposure time of 80 ms. One field per well was recorded for 8 min at 10 frames/s (4800 frames), with two or three wells per treatment in each culture. Wells were imaged sequentially, in an order that alternated between treatment groups. Imaging of the first well started 15 min after medium replacement.

Images were analyzed using the freely available Fluorescence Single Neuron and Network Analysis Package (FluoroSNNAP) [33]. First, cells were automatically segmented in maximum-intensity projection images using Cellpose (v3.1.1.2) [34] with a custom model fine-tuned on images acquired in this experiment (cell diameter 4 pixels, corresponding to approximately 9.5 µm at 2.37 µm/pixel; flow threshold 0.8, cell probability threshold −6). Training labels for fine-tuning were drawn manually by one experimenter. The segmentation masks were then loaded into FluoroSNNAP, in which Ca²⁺ events, transient kinetics, and network synchronization were analyzed with default settings, as described in [33]. Each segmented cell was treated as a region of interest (ROI), and ROIs with at least one detected Ca²⁺ event were classified as active. Because segmentation was based on the somatic fluorescence of Cal-520-loaded cells, ROIs correspond to dye-loaded cell bodies rather than to immunocytochemically identified neurons. Single-cell parameters, including each cell’s maximal participation index and the number of synchronized assemblies in which it participated, were averaged across active ROIs in each well, and well values were then averaged for each treatment within each culture.

### CORT ELISA

CORT concentrations in the plating medium and in medium collected immediately after the fourth medium change, one day before CORT treatment (see section *Medium changes*), were measured with a competitive colorimetric ELISA (Thermo Fisher Scientific, cat. no. EIACORT) according to the manufacturer’s instructions. Because the replacement medium contained no CORT, the latter concentration reflects the baseline level at the onset of treatment. Samples were assayed in duplicate. Absorbance was read at 450 nm within 10 min of adding the stop solution using a Synergy Mx microplate reader (BioTek). CORT concentrations were calculated from a standard curve fitted with a four-parameter logistic (4PL) model and are expressed in nM.

### LDH

Cytotoxicity was assessed by measuring lactate dehydrogenase (LDH) activity in the culture medium with a colorimetric assay (Cytotoxicity Detection Kit^PLUS^ (LDH), Roche, cat. no. 04744926001) according to the manufacturer’s instructions. After 72 h of treatment with CORT (0.25, 0.5, or 1 µM) or vehicle, 50 µl of medium was collected from the upper part of each well to avoid detached cells, transferred to a 96-well plate, and incubated with 100 µl of the reaction mixture for 15 min at room temperature. Absorbance was read at 490 nm using a Synergy Mx microplate reader (BioTek). LDH activity was expressed relative to vehicle-treated controls.

### RNA isolation and RT-qPCR

Total RNA was isolated from primary cortical and hippocampal cultures grown in 6-well plates using the Quick-RNA Miniprep Kit (Zymo Research, cat. no. R1055) according to the manufacturer’s instructions. Cells were lysed in the kit’s RNA lysis buffer, scraped, transferred to tubes, and RNA was eluted in DNase/RNase-free water. RNA concentration and purity were measured with a NanoPhotometer N80 (Implen), and RNA was stored at −80 °C until reverse transcription.

Total RNA (1 µg) was reverse-transcribed into cDNA in a 20-µl reaction using the High-Capacity cDNA Reverse Transcription Kit (Applied Biosystems, cat. no. 4368814) according to the manufacturer’s instructions (25 °C for 10 min, 37 °C for 120 min, 85 °C for 5 min). mRNA expression of special AT-rich sequence-binding protein 2 (*Satb2, Mm00507331_m1*), prospero homeobox 1 (*Prox1, Mm00435969_m1*), FKBP prolyl isomerase 5 (*Fkbp5, Mm00487406_m1*), and beta-2-microglobulin (*B2m, Mm00437762_m1*) was assessed by quantitative real-time PCR (qPCR) using TaqMan Gene Expression Assays (Applied Biosystems). Each 20-µl reaction contained cDNA corresponding to 50 ng of input RNA, 10 µl of TaqMan Fast Advanced Master Mix (Applied Biosystems, cat. no. 4369542), 1 µl of TaqMan assay, and nuclease-free water. Reactions were run in duplicate on a QuantStudio 12K Flex Real-Time PCR System (Applied Biosystems) with the following protocol: UNG incubation at 50 °C for 2 min, polymerase activation at 95 °C for 20 s, and 40 cycles of denaturation at 95 °C for 1 s and annealing/extension at 60 °C for 20 s. Relative expression was calculated with the 2 ^−ΔΔCt^ method [35], using *B2m* as the reference gene, and expressed relative to vehicle (DMSO)-treated controls or, for comparisons between regions, to cortical cultures.

### Statistical analysis

Each experiment was performed in at least three independent biological replicates (except for counting the proportion of cells, where one biological replicate was analyzed), each defined as a separate culture prepared from a single pregnant dam. Measurements from individual dendrites, neurons, images, or wells were first averaged within each biological replicate, and replicate means were used for statistical analysis. The number of biological replicates (n) and the number of dendrites, spines, and neurons analyzed are given in the figure legends. Data were analyzed in a randomized block design, with the replicate as the blocking factor, using paired t-tests for two-group comparisons and one-or two-way repeated-measures (RM) ANOVA for comparisons of more than two groups. Because treatments were randomly assigned to wells within each culture (randomized block design), sphericity was assumed and the Geisser–Greenhouse correction was not applied. To test whether C108 and KET modulated CORT effects differently in cortical and hippocampal neurons, data from both regions were analyzed by three-way mixed ANOVA with region as a between-subject factor and CORT and C108/KET as within-subject factors. Statistical analyses of the relative mRNA levels were performed on log-transformed 2^−ΔΔCt^ values (using ratio paired t-test for two-group comparisons); data are presented as 2^−ΔΔCt^. When values were missing, a mixed-effects model fitted by restricted maximum likelihood (REML) was used instead. A significant effect in one-way RM ANOVA was followed by Dunnett’s post hoc test versus vehicle (DMSO). A significant interaction in two-way RM ANOVA was followed by planned paired t-tests comparing DMSO vs CORT and CORT vs CORT + KET or CORT + C108 and MEM/N2 vs NB/B27. All other comparisons were considered exploratory and were not corrected for multiple testing. No outliers were excluded. Assumptions of parametric tests were verified by inspecting histograms and residual plots. Statistical significance was set at p < 0.05. Full results of all statistical analyses are provided in Supplementary Table S1. Analyses were performed with GraphPad Prism 9.5.1 (GraphPad Software). In graphs, bars represent means, points represent the means of individual biological replicates, and lines connect values from the same replicate across conditions.

### Use of AI tools

Claude (Anthropic) was used for language editing of the manuscript and to assist in writing Python scripts for the selection of representative dendritic segments and representative Ca²⁺ recordings (traces and raster plots), based on their similarity to the group means, and for figure assembly. All scripts and their outputs were verified by the corresponding author, who takes full responsibility for the content of this article.

## Results

### Primary cortical and hippocampal cultures retain regional identity and are predominantly neuronal

To validate the regional identity of the cultures and confirm the accuracy of the dissection, we analyzed the mRNA expression of the region-specific genes *Satb2* and *Prox1* [36]. *Prox1* mRNA, which is typically expressed in hippocampal neurons, was 2.4-fold higher in hippocampal than in cortical cultures (p = 0.039; Supplementary Fig. S1d). *Satb2* mRNA, which is predominantly expressed in cortical neurons, showed the opposite pattern, with approximately 3-fold higher expression in cortical cultures. This difference, however, was not statistically significant (p = 0.086; Supplementary Fig. S1e). To assess the cellular composition of the cultures, we analyzed the proportions of mature neurons (NeuN^+^) and astrocytes (S100B^+^). In a single culture preparation, NeuN^+^ neurons accounted for 58% and 57% of intact nuclei in cortical and hippocampal cultures, respectively, with S100B ^+^ astrocytes representing 6% and 3% (Supplementary Fig. S1a, b, f). Iba1^+^ microglia were virtually absent: in a cortical culture, only 12 Iba1^+^ cells were found in 9 wells (25 fields per well), corresponding to < 0.01% of nuclei (Supplementary Fig. S1c).

### Chronic CORT engages glucocorticoid signaling and reduces dendritic length in a concentration-dependent manner

B27 supplement contains CORT, and because CORT adsorbs to plastic and is degraded over time, its concentration in the medium cannot be predicted [37, 38]. We therefore minimized baseline CORT by successive medium changes and measured its concentration to verify the effectiveness of this protocol. Mean CORT concentration was 21.7 ± 1.8 nM (mean ± SD, n = 3 cultures) in the plating medium and decreased to 0.19 ± 0.18 nM (mean ± SD, n = 3 cultures) after the last medium change, one day before the onset of treatment (Fig. 1a, b).

To verify that CORT engaged glucocorticoid signaling in the cultures, we assessed *Fkbp5* mRNA expression in both cortical and hippocampal cultures. In cortical cultures, chronic CORT increased *Fkbp5* mRNA levels 6-to 8-fold (geometric mean fold change vs DMSO: 7.8, 6.3, and 8.3 for 0.25, 0.5, and 1 µM, respectively; p < 0.0001 vs DMSO for all three concentrations; Fig. 1c). In hippocampal cultures, chronic CORT also increased *Fkbp5* mRNA levels, although to a lesser extent, and only the highest CORT concentration differed significantly from vehicle (geometric mean fold change vs DMSO: 2.4, 2.5, and 4.1 for 0.25, 0.5, and 1 µM, respectively; p = 0.018 vs DMSO for 1 µM; Fig. 1d).

We next evaluated whether chronic CORT influenced cell viability. In cortical cultures, LDH release did not differ from vehicle at any concentration tested (p = 0.146; Fig. 1e), whereas in hippocampal cultures, 1 µM CORT increased LDH release compared with vehicle (p = 0.007; Fig. 1f). We further assessed cell viability by quantifying the number of NeuN^+^ neurons, which did not differ significantly between conditions in either cortical (p = 0.459; Fig. 1g, i) or hippocampal cultures (p = 0.160; Fig. 1h, i). Thus, although 1 µM CORT increased LDH release in hippocampal cultures, the number of NeuN^+^ cells was not reduced, suggesting that the increased LDH release did not reflect substantial neuronal loss.

Finally, we assessed dendritic length after chronic CORT exposure. In cortical cultures, 0.25 µM CORT did not affect total dendritic length, whereas 0.5 and 1 µM CORT decreased it compared with vehicle (p = 0.0007 and p = 0.0002, respectively; Fig. 1j, l). A similar pattern was observed in hippocampal cultures, where 0.25 µM CORT did not differ from vehicle, while 0.5 and 1 µM CORT decreased total dendritic length (p = 0.042 and p = 0.004, respectively; Fig. 1k, l).

Altogether, these data show that chronic CORT engaged glucocorticoid signaling in both cultures, with a stronger transcriptional response in cortical cultures. Cell viability was affected only by 1 µM CORT, and only in hippocampal cultures, whereas 0.5 and 1 µM CORT decreased total dendritic length in both.

### Chronic CORT differentially alters dendritic spine morphology in cortical and hippocampal neurons

To determine how chronic CORT affects dendritic spines, we analyzed spine density and morphology (spine length, head width, length-to-head-width ratio, and spine area) in DiI-labeled cortical and hippocampal neurons after 72 h of exposure to 0.25, 0.5, or 1 µM CORT.

In cortical neurons, chronic CORT produced a nonsignificant trend toward lower spine density (p = 0.097; Fig. 2b), which was not observed at 0.25 µM in subsequent experiments (Figs. 4c, 5b), and did not significantly affect spine length (p = 0.109; Fig. 2c). However, compared with vehicle, all tested CORT concentrations decreased spine head width (p ≤ 0.004 for all concentrations; Fig. 2d) and increased the length-to-head-width ratio (p ≤ 0.002 for all concentrations; Fig. 2e). Spine area was reduced only by 0.5 µM CORT (p = 0.025 vs DMSO; Fig. 2f).

In contrast, in hippocampal neurons, all tested CORT concentrations decreased spine density compared with vehicle (p ≤ 0.048 for all concentrations; Fig. 2h). All CORT concentrations also reduced spine length (p ≤ 0.022 for all concentrations; Fig. 2i) and spine head width (p ≤ 0.005 for all concentrations; Fig. 2j), whereas the length-to-head-width ratio did not differ between conditions (p = 0.173; Fig. 2k). Consistent with the reduction in both spine length and head width, all CORT concentrations decreased spine area compared with vehicle (p ≤ 0.0007 for all concentrations; Fig. 2l). Thus, in cortical neurons, chronic CORT produced spines with narrower heads and a higher length-to-head-width ratio, whereas in hippocampal neurons, spines became proportionally smaller.

The lowest concentration tested (0.25 µM) increased *Fkbp5* expression in cortical cultures (with a nonsignificant 2.4-fold increase in hippocampal cultures) and altered spine morphology in both cortical and hippocampal cultures without affecting cell viability or dendrite length. We therefore used this concentration in all subsequent experiments.

To assess whether culture conditions affect baseline spine morphology, we compared cortical neurons maintained under our protocol (MEM with N2 supplement) with neurons from the same cultures maintained in standard Neurobasal medium with B27 supplement, with or without 0.25 µM CORT. Culture conditions affected spine density (p = 0.0003; Fig. 2n), length (p = 0.001; Fig. 2o), head width (p = 0.002; Fig. 2p), and the length-to-head-width ratio (p = 0.0001; Fig. 2q). Compared with MEM/N2, neurons maintained in Neurobasal/B27 had lower spine density (p = 0.023; Fig. 2n), longer spines (p = 0.049; Fig. 2o) with narrower heads (p = 0.015; Fig. 2p), and a higher length-to-head-width ratio (p = 0.002; Fig. 2q). Spine area, in contrast, did not differ between groups (p = 0.613; Fig. 2r). In Neurobasal/B27, 0.25 µM CORT did not further change any of these parameters (all p > 0.06). Because the two conditions differed in both the basal medium and the supplement, these differences cannot be attributed to CORT alone; however, the narrower spine heads and higher length-to-head-width ratio in Neurobasal/B27 resembled the effects of chronic CORT under our culture conditions.

### Chronic CORT does not affect PSD-95 puncta in cortical or hippocampal neurons

Spine size correlates with synapse size and stability, as well as with the abundance of postsynaptic scaffold proteins [39, 40]. Among these proteins, PSD-95, the major scaffold of the postsynaptic density, is a regulator of synapse size and stability [41]. Thus, to determine whether chronic CORT-induced changes in spine morphology are accompanied by alterations in PSD-95 abundance, we quantified PSD-95 immunoreactivity within masks of DiI-labeled dendrites and their spines. In cortical neurons, chronic CORT (0.25 µM) did not significantly change PSD-95 puncta density (p = 0.340; Fig. 3b), puncta size (p = 0.555; Fig. 3c), or mean PSD-95 fluorescence intensity (p = 0.353; Fig. 3d). Similarly, in hippocampal neurons, PSD-95 puncta density (p = 0.263; Fig. 3f), puncta size (p = 0.859; Fig. 3g), and mean PSD-95 fluorescence intensity (p = 0.792; Fig. 3h) did not differ from vehicle. Thus, CORT-induced changes in spine morphology occurred without detectable changes in dendritic PSD-95.

**Fig. 3.**
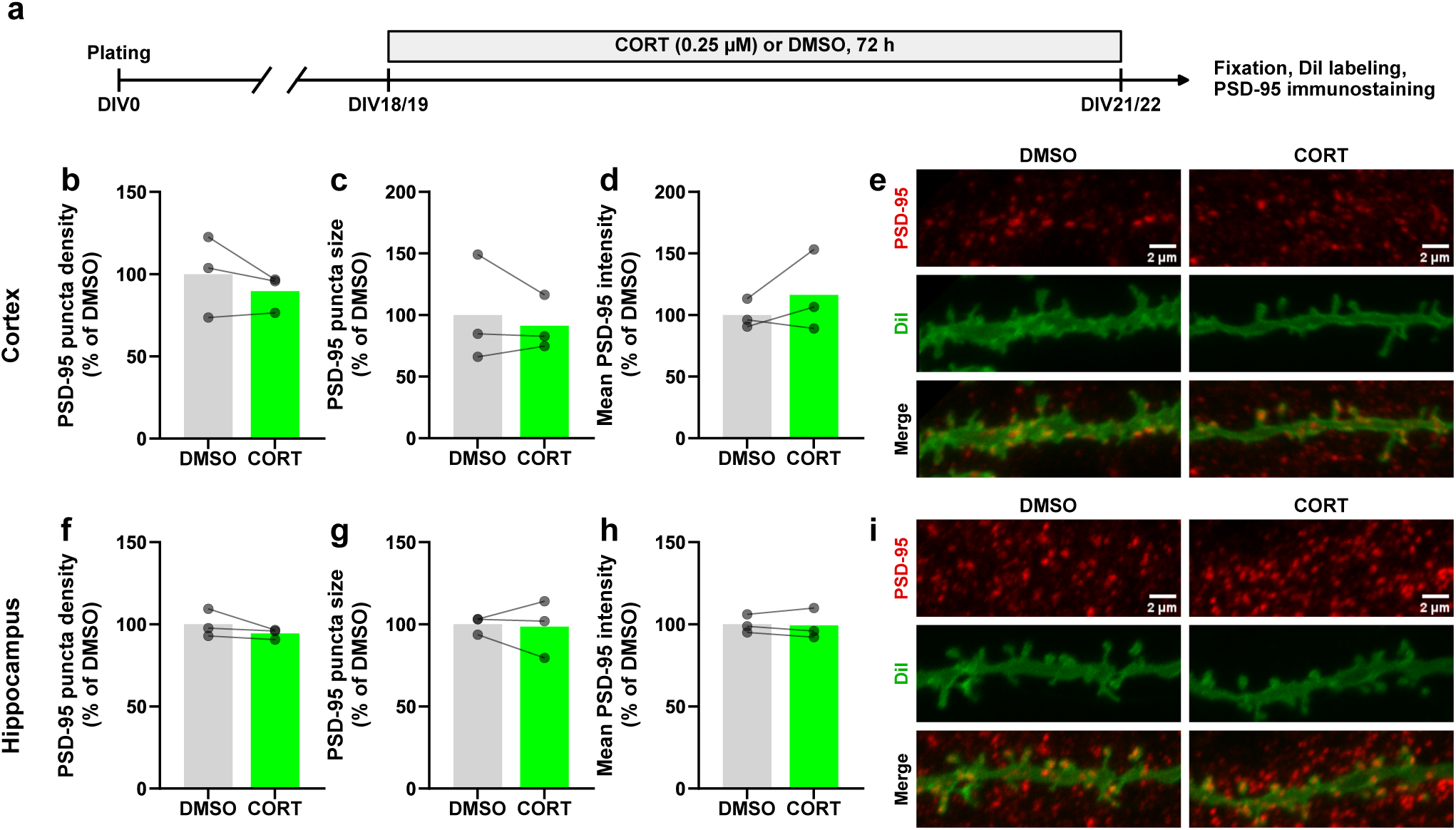
Chronic CORT does not change dendritic PSD-95 in cortical or hippocampal neurons. **a** Experimental timeline. Cultures were treated with CORT (0.25 µM) or DMSO for 72 h from DIV18/19, fixed at DIV21/22, labeled with DiI, and immunostained for PSD-95. **b–d** Density (**b**), size (**c**) of PSD-95 puncta and mean PSD-95 fluorescence intensity (**d**) within DiI-labeled dendrites and spines of cortical neurons, expressed as % of DMSO. **e** Representative images of PSD-95 immunostaining (red), DiI-labeled dendrites (green), and merged images from cortical cultures. **f–i** The same parameters and images for hippocampal neurons. In total, 68 and 75 dendrites were analyzed in cortical and hippocampal cultures, respectively, n = 3 cultures; none of the comparisons was significant (paired t-test). Bars represent means, points represent individual cultures, and lines connect values from the same culture. Scale bar: 2 µm

### Selective GR modulation attenuates CORT effects on spine head width in cortical neurons

To determine whether the effects of chronic CORT on dendritic spine morphology were mediated by GR, we co-treated cultures with CORT and the selective GR modulator C108. To assess whether C108 interfered with GR-dependent transcription at the concentration used, we first measured mRNA levels of the canonical GR target gene *Fkbp5* in cortical cultures.

CORT increased *Fkbp5* mRNA levels (p = 0.0007; Fig. 4b). The CORT-induced increase was smaller in the presence of C108 (geometric mean fold change: 5.4 for CORT vs DMSO and 2.1 for CORT + C108 vs C108); however, the CORT × C108 interaction did not reach significance (p = 0.085; Fig. 4b).

**Fig. 4.**
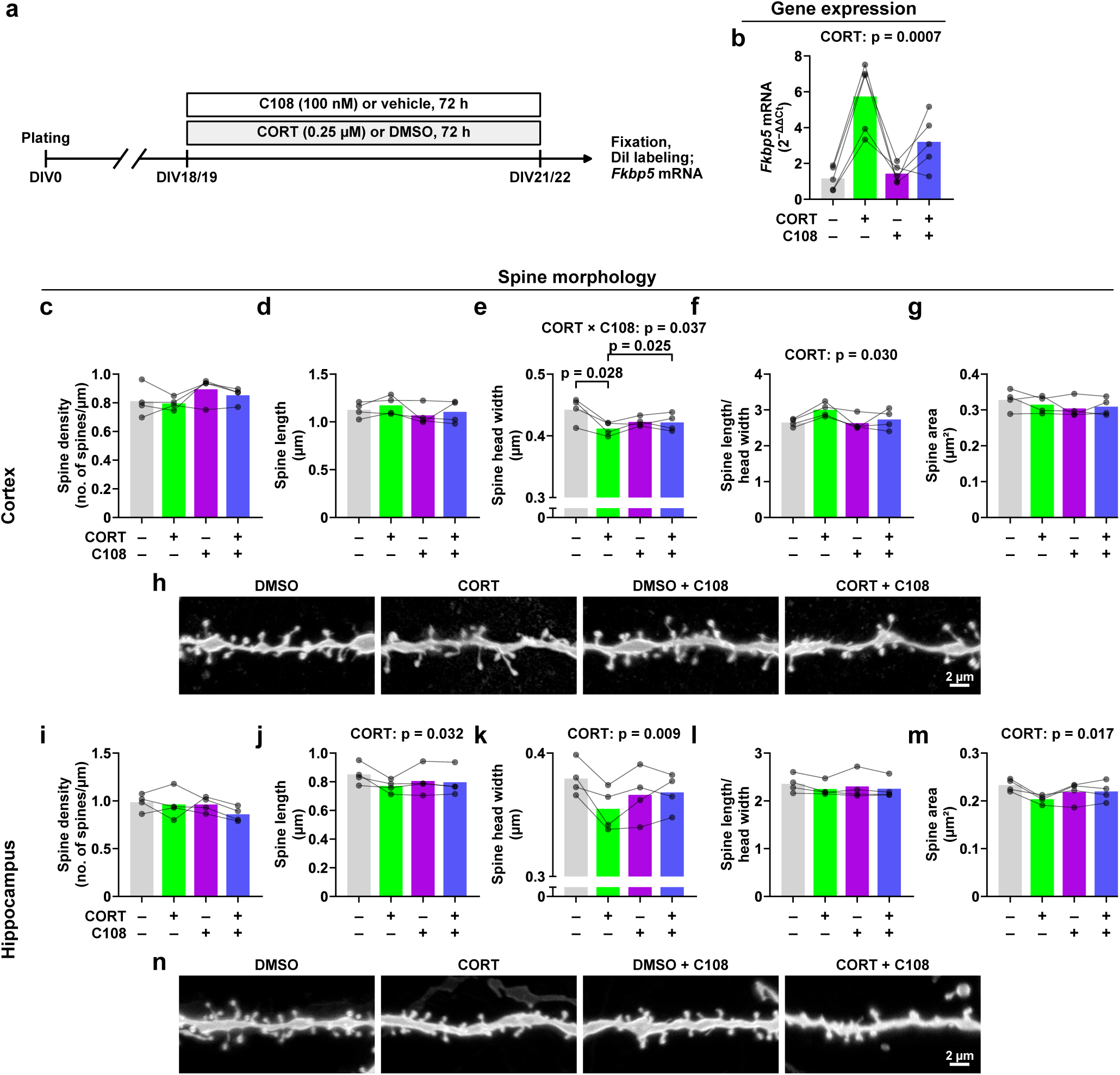
The selective GR modulator C108 prevents the CORT-induced decrease in spine head width in cortical neurons. **a** Experimental timeline. Cultures were treated with CORT (0.25 µM) or DMSO and with C108 (100 nM) or vehicle for 72 h from DIV18/19; spines were analyzed and *Fkbp5* mRNA was measured at DIV21/22. **b** *Fkbp5* mRNA levels in cortical cultures relative to DMSO (n = 5 cultures). Bars in **b** represent arithmetic means of 2^−ΔΔCt^ values, whereas statistical analyses were performed on log-transformed values and geometric mean fold changes are given in the text. **c–g** Spine density (**c**), spine length (**d**), spine head width (**e**), length-to-head-width ratio (**f**), and spine area (**g**) in cortical neurons. **h** Representative DiI-labeled dendritic segments (20 µm) of cortical neurons. **i–m** The same parameters in hippocampal neurons. **n** Representative dendritic segments of hippocampal neurons. In total, 9916 spines on 12,049 µm of dendrites were analyzed in cortical cultures and 8923 spines on 9671 µm of dendrites in hippocampal cultures; n = 4 cultures. Bars represent means, points represent individual cultures, and lines connect values from the same culture. Significant main effects and interactions from two-way RM ANOVA are shown above the graphs; for a significant interaction, p values from planned paired t-tests (DMSO vs CORT; CORT vs CORT + C108) are shown. In **h** and **n**, segments were selected from one culture so that the local spine density, length, head width, and area within the segment approximated the group means and preserved the direction of differences between groups. Scale bars: 2 µm

In cortical neurons, C108 prevented the CORT-induced decrease in spine head width (CORT × C108 interaction: p = 0.037; Fig. 4e). CORT decreased spine head width (DMSO vs CORT: p = 0.028; Fig. 4e), and spine heads were wider in the CORT + C108 group than in the CORT group (CORT vs CORT + C108: p = 0.025; Fig. 4e). In exploratory comparisons, CORT did not affect spine head width in the presence of C108 (C108 vs CORT + C108: p = 0.866; Fig. 4e), and C108 alone produced only a nonsignificant trend toward narrower spine heads (DMSO vs C108: p = 0.084; Fig. 4e). CORT also increased the length-to-head-width ratio, and this effect was not modified by C108 (CORT: p = 0.030; CORT × C108: p = 0.293; Fig. 4f).

In hippocampal neurons, CORT reduced spine length (p = 0.032; Fig. 4j), head width (p = 0.009; Fig. 4k), and area (p = 0.017; Fig. 4m). C108 attenuated these effects to a similar extent as in cortical neurons (CORT effect on spine head width: −6.5% without and +0.5% with C108; spine area: −12.7% and +0.6%), although the CORT × C108 interactions did not reach significance. The modulation of CORT effects by C108 did not differ between regions (region × CORT × C108 interaction: all p > 0.16).

Neither CORT nor C108 affected the remaining parameters, namely spine density in cortical and hippocampal neurons, spine length and area in cortical neurons, and the length-to-head-width ratio in hippocampal neurons (all p ≥ 0.077; Fig. 4c, d, g, i, l).

These results indicate that GR mediates the effect of chronic CORT on spine head width in cortical neurons. In hippocampal neurons, C108 attenuated CORT effects to a similar extent, but GR dependence could not be established statistically.

### Ketamine reverses CORT-induced reductions in spine size in hippocampal neurons

To determine whether KET counteracts the effects of chronic CORT on spine morphology, KET (10 µM) was added to CORT-or vehicle-treated cultures during the last 24 h of treatment.

In cortical neurons, KET increased spine length, whereas CORT did not affect it (KET: p = 0.024; CORT: p > 0.99; Fig. 5c). CORT decreased spine head width regardless of KET (CORT: p = 0.042; CORT × KET: p = 0.139; Fig. 5d), whereas its effect on the length-to-head-width ratio was modified by KET (CORT × KET: p = 0.018; Fig. 5e). CORT increased this ratio (DMSO vs CORT: p = 0.012; Fig. 5e); however, CORT + KET did not differ significantly from CORT (CORT vs CORT + KET: p = 0.109; Fig. 5e). Exploratory analysis showed that KET alone also increased the length-to-head-width ratio (DMSO vs KET: p = 0.010; Fig. 5e).

**Fig. 5.**
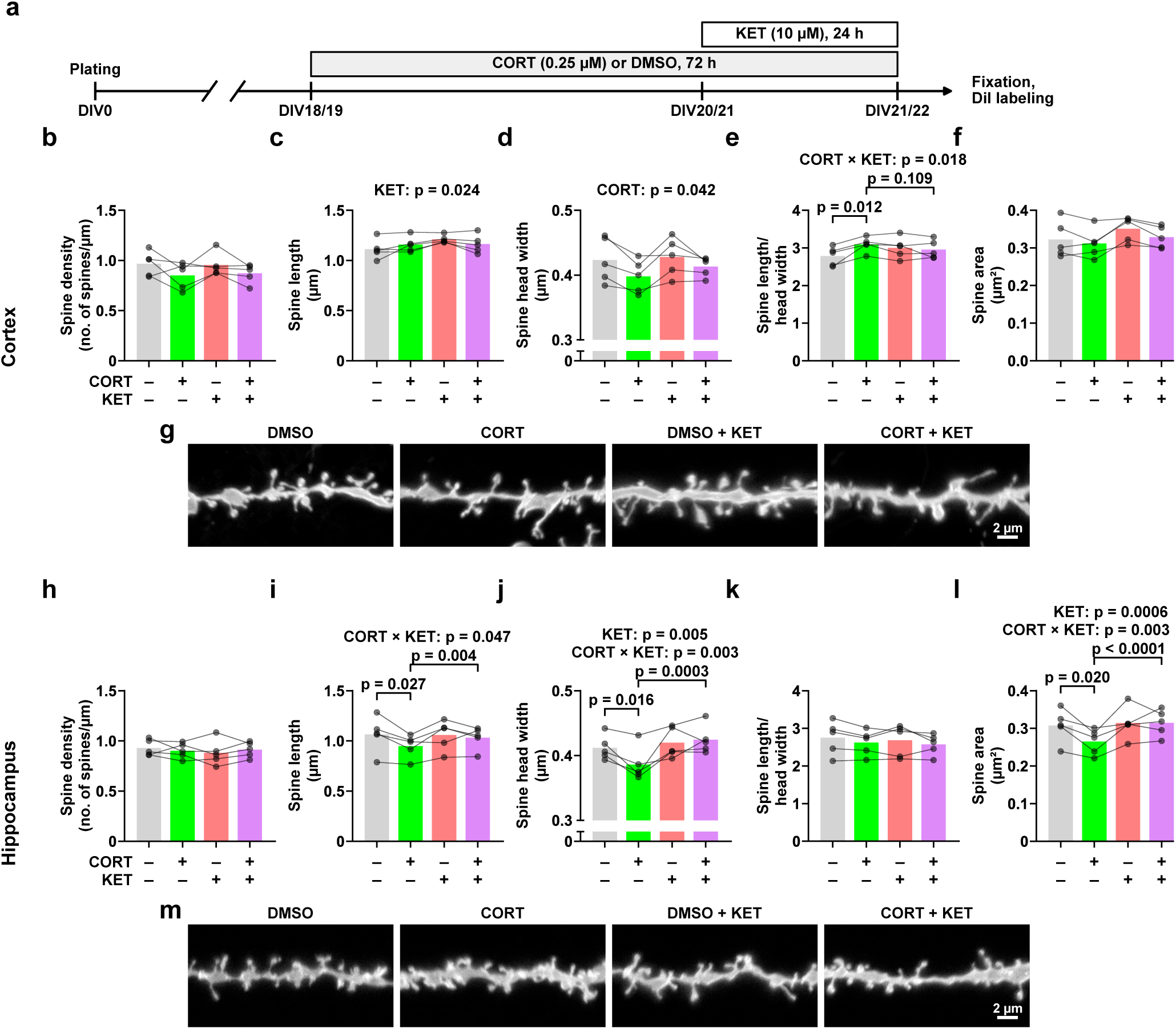
KET reverses the CORT-induced reduction in spine size in hippocampal but not cortical neurons. **a** Experimental timeline. Cultures were treated with CORT (0.25 µM) or DMSO for 72 h from DIV18/19, with or without KET (10 µM) for the last 24 h, and fixed and labeled with DiI at DIV21/22. **b–f** Spine density (**b**), spine length (**c**), spine head width (**d**), length-to-head-width ratio (**e**), and spine area (**f**) in cortical neurons. **g** Representative DiI-labeled dendritic segments (20 µm) of cortical neurons. **h–l** The same parameters in hippocampal neurons. **m** Representative dendritic segments of hippocampal neurons. In total, 14,029 spines on 15,652 µm of dendrites were analyzed in cortical cultures and 17,358 spines on 19,152 µm of dendrites in hippocampal cultures; n = 5 cultures. Bars represent means, points represent individual cultures, and lines connect values from the same culture. Significant main effects and interactions from two-way RM ANOVA are shown above the graphs; for a significant interaction, p values from planned paired t-tests (DMSO vs CORT; CORT vs CORT + KET) are shown. In **g** and **m**, segments were selected as in Fig. 4. Scale bars: 2 µm

In hippocampal neurons, KET reversed the CORT-induced reduction in spine size. The CORT × KET interaction was significant for spine length (p = 0.047; Fig. 5i), head width (p = 0.003; Fig. 5j), and area (p = 0.003; Fig. 5l). CORT decreased spine length (DMSO vs CORT: p = 0.027; Fig. 5i), head width (p = 0.016; Fig. 5j), and area (p = 0.020; Fig. 5l), and all three parameters were higher in the CORT + KET group than in the CORT group (CORT vs CORT + KET: p = 0.004, p = 0.0003, and p < 0.0001, respectively; Fig. 5i, j, l). In exploratory comparisons, KET alone did not affect spine length, head width, or area (DMSO vs KET: all p > 0.26), and spine length and area in the CORT + KET group did not differ from DMSO (p = 0.441 and p = 0.513, respectively), whereas spine head width was slightly larger than in DMSO (p = 0.028).

Neither CORT nor KET affected the remaining parameters, namely spine density in cortical and hippocampal neurons, spine area in cortical neurons, and the length-to-head-width ratio in hippocampal neurons (all p ≥ 0.088; Fig. 5b, f, h, k).

The modulation of CORT effects by KET differed between regions (region × CORT × KET interaction: p ≤ 0.031 for spine head width, length, area, and length-to-head-width ratio).

Thus, in cortical neurons, KET increased spine length but did not counteract the effects of CORT, whereas in hippocampal neurons, KET reversed the CORT-induced reduction in spine length, head width, and area, although KET alone had no effect on these parameters.

### Chronic CORT alters neuronal network activity in a region-specific manner, whereas ketamine reduces network synchronization

To determine whether the structural effects of CORT and KET are accompanied by changes in cell function, we measured spontaneous Ca²⁺ activity in cortical and hippocampal cultures at the end of treatment (DIV21 or 22; 8 min at 10 Hz), in the continued presence of vehicle, CORT, and/or KET (Figs. 6 and 7). All analyses were performed on culture means.

**Fig. 6.**
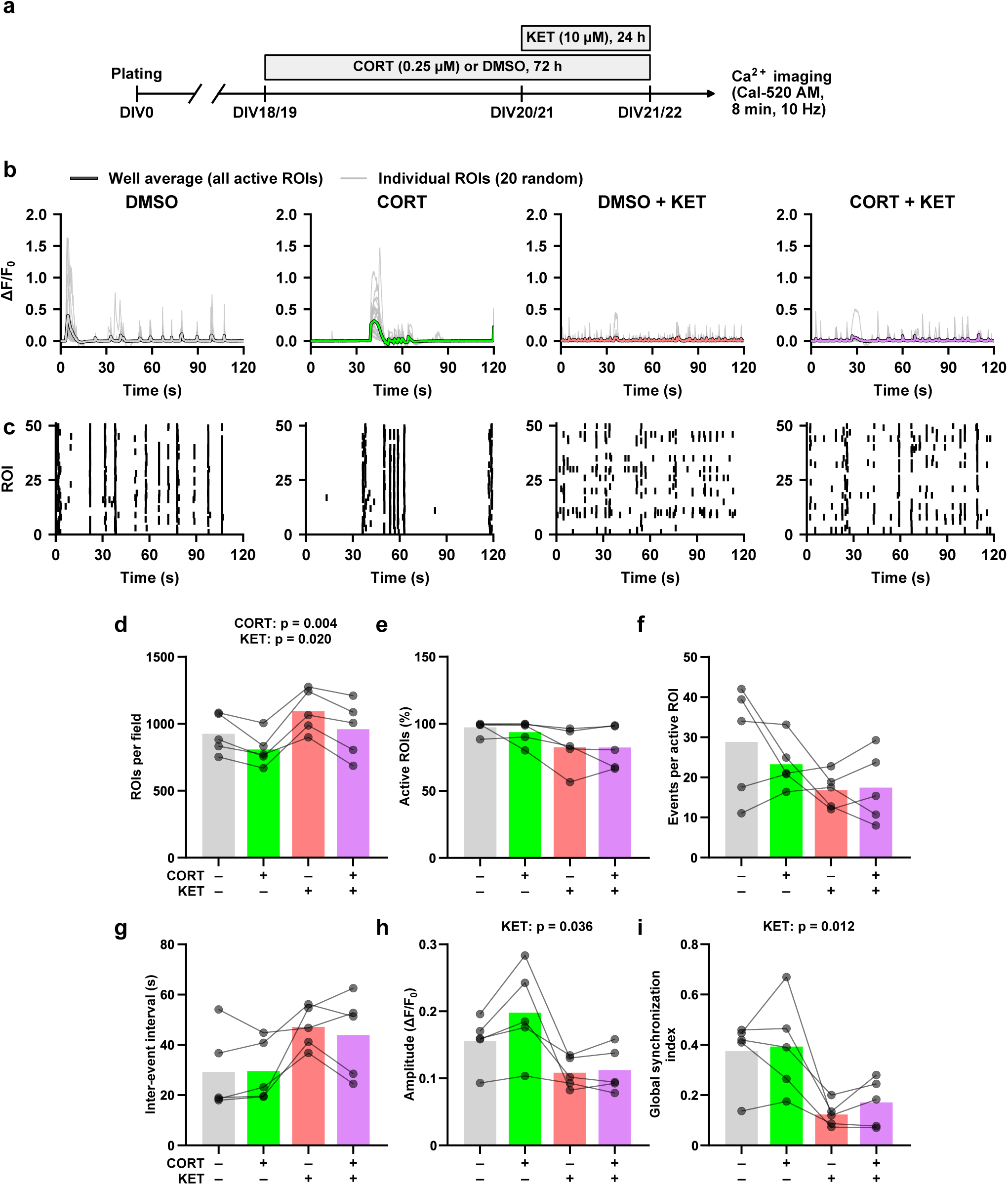
CORT and KET alter spontaneous Ca²⁺ activity in cortical cultures. **a** Experimental timeline. Cultures were treated with CORT (0.25 µM) or DMSO for 72 h from DIV18/19, with or without KET (10 µM) for the last 24 h. Spontaneous Ca²⁺ activity was recorded at DIV21/22 with Cal-520 AM (8 min, 10 Hz) in the continued presence of the drugs. **b** Representative ΔF/F₀ traces (120-s windows): mean of all active ROIs in the well (colored line) and 20 randomly selected active ROIs (gray). **c** Raster plots of Ca²⁺ event onsets in 50 active ROIs from the same windows. For each condition, the well and 120-s window closest to the group means of the number of events per active ROI, amplitude, and global synchronization index were selected from one culture. **d–i** Number of ROIs per field (**d**), proportion of active ROIs (≥ 1 detected event; **e**), number of events per active ROI (**f**), inter-event interval (**g**), amplitude of Ca²⁺ transients (**h**), and global synchronization index (**i**). Values were averaged across wells (two or three per condition) within each culture; n = 5 cultures. Bars represent means, points represent individual cultures, and lines connect values from the same culture. Significant main effects and interactions from two-way RM ANOVA are shown above the graphs

**Fig. 7.**
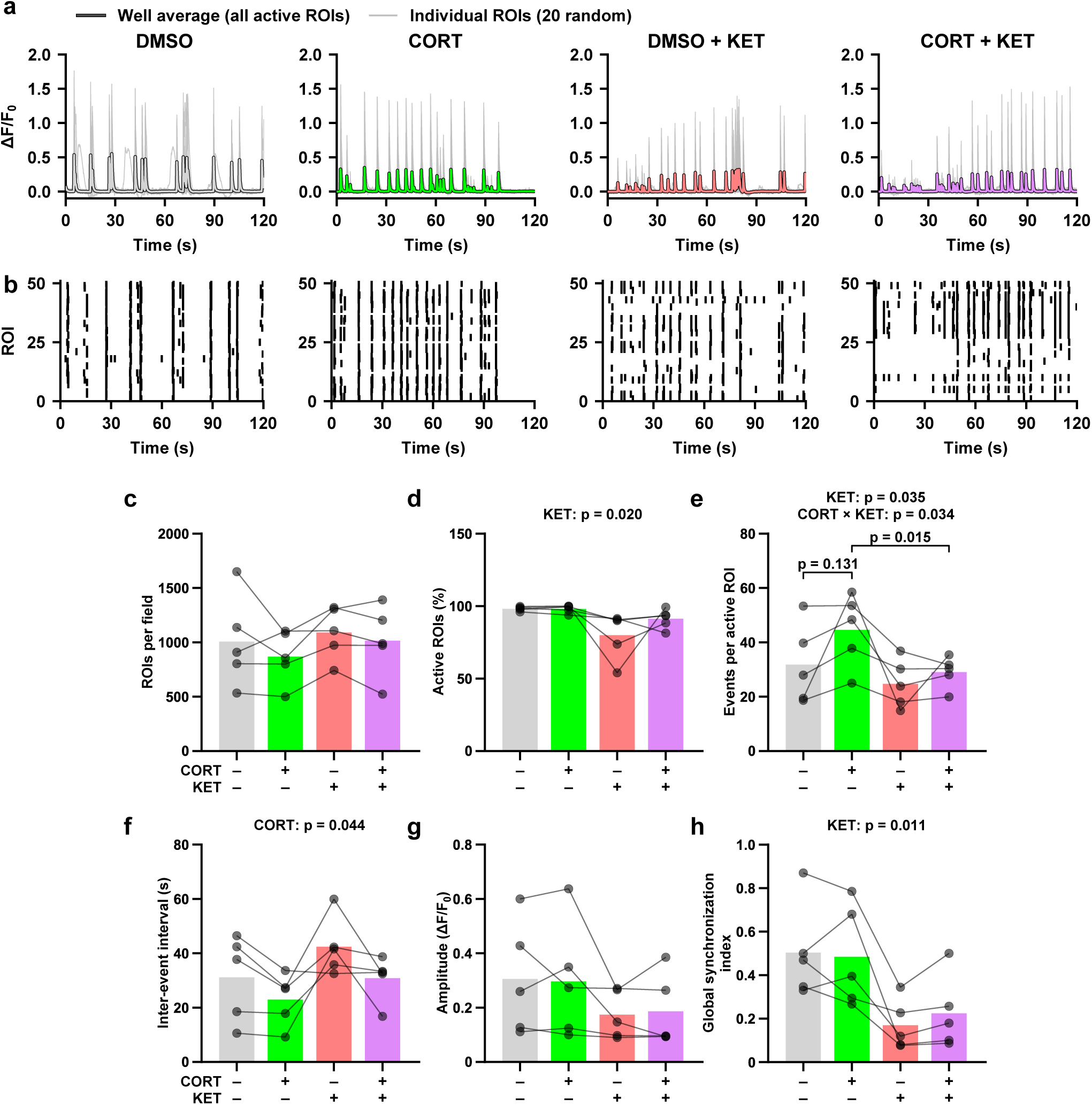
CORT and KET alter spontaneous Ca²⁺ activity in hippocampal cultures. Treatment and recording as in Fig. 6a. **a** Representative ΔF/F₀ traces (120-s windows): mean of all active ROIs in the well (colored line) and 20 randomly selected active ROIs (gray). **b** Raster plots of Ca²⁺ event onsets in 50 active ROIs from the same windows. Wells and windows were selected as in Fig. 6. **c–h** Number of ROIs per field (**c**), proportion of active ROIs (≥ 1 detected event; **d**), number of events per active ROI (**e**), inter-event interval (**f**), amplitude of Ca²⁺ transients (**g**), and global synchronization index (**h**); n = 5 cultures. Bars represent means, points represent individual cultures, and lines connect values from the same culture. Significant main effects and interactions from two-way RM ANOVA are shown above the graphs; for a significant interaction, p values from planned paired t-tests (DMSO vs CORT; CORT vs CORT + KET) are shown

In cortical cultures, CORT reduced the number of ROIs (segmented cells) per field, which KET increased (CORT: p = 0.004; KET: p = 0.020; Fig. 6d). This decrease was not due to cell loss, as 0.25 µM CORT affected neither LDH release nor the number of NeuN⁺ cells (Fig. 1e–h), and the proportion of active ROIs was unchanged. CORT also reduced the number of synchronized assemblies per cell (p = 0.013; Supplementary Fig. S2a). CORT did not affect the other parameters (all p > 0.35), apart from a nonsignificant trend toward larger transient amplitude (p = 0.068; Fig. 6h).

KET decreased global synchronization (p = 0.012; Fig. 6i) and transient amplitude (p = 0.036; Fig. 6h), whereas its effect on the number of events per active ROI did not reach significance (p = 0.094; Fig. 6f). KET also showed nonsignificant trends toward a lower proportion of active ROIs (p = 0.054; Fig. 6e) and longer inter-event intervals (p = 0.070; Fig. 6g), with no CORT × KET interactions (all p > 0.37). Among the additional parameters (Supplementary Fig. S2), KET decreased the participation index, which is derived from the same synchronization analysis as the global synchronization index (p = 0.019; Supplementary Fig. S2b), and the coefficient of variation of transient amplitude (p = 0.010; Supplementary Fig. S2c), whereas neither treatment affected the rise or fall times of Ca²⁺ transients (all p ≥ 0.055; Supplementary Fig. S2d, e).

In hippocampal cultures, the effect of CORT on the number of events per active ROI was modified by KET (CORT × KET: p = 0.034; Fig. 7e): CORT increased this number by 40%, although not significantly (DMSO vs CORT: p = 0.131; Fig. 7e), and KET reduced it in CORT-treated cultures (CORT vs CORT + KET: p = 0.015; Fig. 7e), so that CORT + KET did not differ from vehicle (exploratory comparison, DMSO vs CORT + KET: p = 0.694; Fig. 7e). Consistent with increased event frequency, CORT shortened the inter-event interval (p = 0.044; Fig. 7f). Neither treatment affected the number of ROIs per field (all p ≥ 0.18; Fig. 7c), and CORT did not affect the other parameters (all p ≥ 0.26), apart from the rise time of Ca²⁺ transients (see below).

KET decreased global synchronization (p = 0.011; Fig. 7h) and the proportion of active ROIs (p = 0.020; Fig. 7d), whereas its effect on transient amplitude did not reach significance (p = 0.082; Fig. 7g). Among the additional parameters (Supplementary Fig. S2), the effect of KET on the number of assemblies per cell did not reach significance (p = 0.050; Supplementary Fig. S2f), whereas KET decreased the participation index (p = 0.012; Supplementary Fig. S2g). Neither treatment affected the coefficient of variation of transient amplitude (all p > 0.14; Supplementary Fig. S2h). The rise time of Ca²⁺ transients was affected by CORT, KET, and their interaction (CORT: p = 0.005; KET: p = 0.016; CORT × KET: p = 0.017; Supplementary Fig. S2i), and the CORT × KET interaction was also significant for fall time (CORT × KET: p = 0.040; Supplementary Fig. S2j); however, neither parameter differed between DMSO and CORT or between CORT and CORT + KET (all p ≥ 0.16; Supplementary Fig. S2i, j).

Thus, CORT shortened inter-event intervals in hippocampal but not in cortical cultures, whereas KET reduced network synchronization in both regions, independently of CORT.

## Discussion

Prolonged exposure to CORT is known to reduce the density of dendritic spines in the prefrontal cortex and hippocampus [11, 16]. However, much less is known about how chronic CORT affects spine morphology and whether these effects differ between cortical and hippocampal neurons. It is also unknown whether KET, which restores spines eliminated by chronic CORT in the prefrontal cortex [12], counteracts these morphological changes, and whether it does so in cultures from both regions. Here, we show that chronic CORT, at a concentration that increased *Fkbp5* expression without affecting cell viability or dendrite length, altered spine morphology in a region-specific manner: cortical neurons developed spines with narrower heads and a higher length-to-head-width ratio, with the decrease in spine head width depending on GR, whereas hippocampal neurons developed proportionally smaller spines. KET reversed these structural changes in hippocampal but not in cortical neurons. To relate these changes to function, we also recorded spontaneous Ca²⁺ activity, and CORT effects on network activity were likewise region-specific, with Ca²⁺ events becoming more frequent in hippocampal but not in cortical cultures. We propose that CORT reshapes spines by two different routes in the two cultures. In cortical neurons, the narrowing of spine heads was GR-dependent and occurred without an increase in network activity. In hippocampal neurons, spine shrinkage may be at least partly secondary to this increase in activity. Although CORT likely acts through GR in both cultures, it is this difference in network context that may determine why KET reversed the structural changes only in hippocampal neurons.

We used primary neuronal cultures to obtain a reduced experimental system in which the effects of CORT on neuronal structure could be assessed more directly, with less influence from the many interacting factors present in the intact brain. Because there is no standardized protocol for CORT treatment of primary neurons, we selected the lowest tested concentration that increased expression of the GR target gene *Fkbp5* in cortical cultures (with a similar, nonsignificant trend in hippocampal cultures) and altered spine morphology in both cultures without affecting cell viability or total dendritic length. Although relatively high, this concentration is comparable to CORT levels measured in rat brain tissue after acute stress (≈ 275 nM) [13] and within the range commonly used in vitro [28, 38]. Concentrations above physiological levels are generally required in vitro, because neurons are maintained in B27, which already contains CORT, and because steroids adsorb to culture plasticware, so that the effective concentration may be lower than the nominal one [38]. For the same reasons, the CORT concentration in B27 cannot be predicted; we therefore reduced it to sub-nanomolar levels through successive medium changes. This step was relevant for spine morphology: cortical neurons maintained in standard Neurobasal/B27 medium had narrower spine heads and a higher length-to-head-width ratio, resembling the effects of chronic CORT, and 0.25 µM CORT did not further alter spine morphology in these conditions. Although differences between the two media cannot be attributed to CORT alone, these results suggest that standard B27-based conditions may partially mask CORT effects on dendritic spines.

Under these conditions, we first compared our data with in vivo findings, which constitute most of the available literature on CORT effects on dendritic spines [7]. These studies, however, concern mainly spine density and turnover or describe morphology either by single size measures or by classifying spines into subtypes using fixed thresholds, although spine shapes form a continuum [12, 16, 42]. Moreover, they are derived from systemic CORT administration, which makes it difficult to distinguish direct effects of CORT on neurons from indirect ones [15, 43]. In many studies, chronic CORT produced region-specific changes, reducing spine density in the prefrontal cortex and hippocampus and increasing it in the amygdala [11, 16]. In contrast to these reports, in our studies CORT did not affect spine density in cortical neurons, and in hippocampal neurons, a reduction was observed in one experiment but not in two others. These differences may reflect indirect effects of CORT that are absent in vitro. In particular, stress-induced spine loss in vivo involves microglia-mediated synaptic elimination [44, 45], which can depend on GR [46], whereas Iba1^+^ microglia were virtually absent from our cultures (Supplementary Fig. S1c). In addition, the shorter exposure (72 h) may have limited spine loss. Thus, in the absence of systemic and microglial influences, the effects of CORT primarily concerned spine morphology rather than spine number.

Because the effects of CORT in our cultures concerned mainly spine morphology, their comparison with previous studies is limited, as quantitative data on CORT-induced changes in spine morphology are scarce, particularly in vitro. To our knowledge, only one in vitro study has addressed this question: in primary rat hippocampal neurons maintained in CORT-free B27, CORT reduced spine head size, decreased the proportion of mushroom spines, and increased the proportion of thin spines, with effects observed from 100 nM [13]. These changes are consistent with our hippocampal data, in which spines became shorter and their heads narrower, with an unchanged length-to-head-width ratio. Similar data for cortical neurons in vitro are lacking. In our cortical cultures, CORT narrowed spine heads without changing spine length, thereby increasing the length-to-head-width ratio, a pattern suggestive of a shift toward filopodia-like morphology. In vivo, chronic CORT or stress reduced spine density in several prefrontal cortex regions [11, 16], with subtype-selective loss that differs between studies [16, 47] and an increased proportion of small-headed spines [14], which is consistent with the narrowing of spine heads in our cortical cultures. Thus, although CORT affected different morphological parameters in cortical and hippocampal neurons, in both regions the changes are consistent with a shift toward a less mature spine phenotype.

However, spine shape alone does not reveal whether these changes affect synapse stability. Spine head volume correlates with PSD size [48, 49], and loss of PSD-95, a major PSD scaffolding protein, increases the likelihood of spine elimination [50, 51]. Yet the morphological changes induced by CORT were not accompanied by detectable changes in PSD-95 puncta density, size, or intensity in either cortical or hippocampal neurons. Because PSD-95 was quantified in masks including both dendritic shafts and spines, a decrease restricted to spines may have been diluted by the shaft signal, particularly since cultured neurons form a larger fraction of excitatory synapses on dendritic shafts than neurons in vivo [52–54], and spine retraction can relocate PSD-95 to the shaft. Moreover, the synaptic function of PSD-95 is regulated by activity-dependent palmitoylation [55, 56], which was not assessed here and may change without alterations in PSD-95 abundance. Nevertheless, the lack of CORT effects on PSD-95 is consistent with previous studies in which CORT altered neuronal excitability without affecting PSD-95 clustering or expression [57, 58]. This suggests that, rather than through changes in the postsynaptic scaffold, CORT may act on synapses primarily through changes in synaptic transmission and network activity.

Indeed, structural and functional plasticity are bidirectionally linked: changes in network activity remodel dendritic spines, and spine morphology, in turn, determines synaptic strength and thereby network excitability [4, 59, 60]. In our cortical cultures, CORT did not change the frequency of Ca²⁺ events or network synchronization but tended to increase the amplitude of Ca²⁺ transients. This may be related to previous electrophysiological reports in which brief CORT exposure increased AMPAR-and NMDAR-mediated transmission in a GR-dependent and sustained manner in prefrontal cortex slices, and increased whole-cell AMPAR and NMDAR current density and miniature excitatory postsynaptic current (mEPSC) amplitude without changing mEPSC frequency in cultured prefrontal neurons [57]. In contrast, repeated stress in vivo and prolonged (5–7 day) CORT exposure in prefrontal cultures suppress glutamate receptor expression and AMPAR-mediated synaptic transmission through GR [58]. Although Ca²⁺ transient amplitude does not directly reflect synaptic currents, the tendency toward larger transients without a change in event frequency after 72 h of CORT resembles the effects of brief rather than prolonged glucocorticoid exposure. Together with the GR dependence of spine head narrowing, these findings are consistent with a postsynaptic, GR-mediated action of CORT in cortical neurons. The GR dependence was, however, statistically established only for spine head width. This may reflect incomplete GR blockade: at the concentration used, C108 only partially attenuated CORT-induced *Fkbp5* expression. Moreover, C108 alone tended to reduce spine head width, mimicking the effect of CORT, consistent with partial agonist activity at GR [27], so that values in the CORT + C108 group were closer to those in the C108 group than to those in the vehicle group. Such residual GR activation may therefore have prevented the detection of antagonism for the CORT-induced increase in the length-to-head-width ratio.

In our hippocampal cultures, by contrast, CORT did not affect the amplitude but tended to increase the frequency of Ca²⁺ events (by 40%) and shortened the inter-event interval. In the hippocampus, corticosteroids rapidly increase mEPSC frequency by enhancing glutamate release through membrane MR [61]. Prolonged network hyperactivity triggers homeostatic downscaling of excitatory synapses, which is accompanied by spine shrinkage [62, 63]. Because Ca²⁺ events occurred more frequently in our CORT-treated hippocampal cultures, the reduction in spine size may therefore be at least partly a consequence of CORT-induced hyperactivity. Such hyperactivity may involve MR, which is highly expressed in the hippocampus [8, 64], would be fully occupied at the CORT concentration used here, and is not targeted by C108. This does not, however, exclude a contribution of GR. In hippocampal neurons, C108 attenuated the effects of CORT on spine head width and area to a similar extent as in cortical neurons, and the modulation of CORT effects by C108 did not differ between cultures. Nevertheless, the CORT × C108 interactions did not reach significance, which may reflect variability between cultures. As in cortical neurons, C108 alone also tended to reduce these parameters.

This contribution of network activity to CORT-induced changes in the hippocampus may also explain the region-specific effects of KET. In hippocampal cultures, KET reduced the number of Ca²⁺ events per active cell in CORT-treated cultures to control levels and, in parallel, reversed CORT-induced changes in spine size. By reducing CORT-induced hyperactivity, KET may have removed the drive for homeostatic downscaling of hippocampal synapses. Moreover, by blocking NMDAR activity at rest, KET inhibits eukaryotic elongation factor 2 kinase and thereby rapidly increases BDNF synthesis [65], triggering homeostatic synaptic upscaling [66], which may counteract the downscaling-like reduction in spine size induced by CORT. Because NMDAR blockade at rest does not require elevated network activity, this mechanism would be expected to operate in cortical neurons as well. In these neurons, however, KET did not counteract the GR-dependent narrowing of spine heads, possibly because, unlike the activity-dependent trigger of spine shrinkage in hippocampal neurons, GR activation persisted during KET treatment. Instead, KET independently increased spine length. This elongation resembles the shift toward thin and filopodial spines induced by serotonergic psychedelics in cortical cultures, although, unlike psychedelics, KET did not increase spine density [25]. In vivo, KET protects mushroom spines in the prefrontal cortex from stress-induced elimination potentially through activation of parvalbumin-expressing interneurons [67] and restores spines lost after chronic CORT or stress within 12–24 h [12]. In our cultures, however, CORT did not reduce cortical spine density, leaving no spine loss for KET to reverse.

Independently of CORT, KET reduced network synchronization in both regions. In culture, NMDARs contribute substantially to synchronized network bursts, mainly by sustaining them after AMPAR-driven initiation [68], and KET reduces spiking in cortical cultures already at 500 nM and, at 10 µM, also network bursting [69]. In vivo, KET similarly decreases Ca²⁺ event rates in hippocampal CA1 [70], whereas in the neocortex it does not globally suppress activity but switches it between neuronal populations, silencing previously active pyramidal neurons and activating previously silent ones through interneuron-mediated disinhibition [71]. Because Ca²⁺ activity was recorded in the continued presence of KET, the functional effects in our cultures likely reflect, at least in part, acute NMDAR blockade, and differences from in vivo findings may also arise from the absence of afferent and neuromodulatory inputs. Whether the KET-induced structural rescue in hippocampal neurons translates into lasting functional normalization therefore remains to be determined.

In conclusion, our results show that, in the absence of systemic influences and in cultures containing low proportions of mature astrocytes (S100B⁺) and microglia (Iba1⁺), chronic CORT primarily alters spine morphology rather than spine number, and that these effects, as well as their reversibility by KET, are region-specific. In cortical neurons, CORT narrowed spine heads through GR without changing the frequency of Ca²⁺ events, and this change was not counteracted by KET. In hippocampal neurons, CORT shortened the intervals between Ca²⁺ events, accompanied by a nonsignificant 40% increase in the number of events per active cell, and reduced spine size, possibly through homeostatic mechanisms. KET reversed the reduction in spine size and reduced the number of Ca²⁺ events per active cell in CORT-treated cultures to control levels. These findings suggest that the efficacy of rapid-acting antidepressants in counteracting stress-related synaptic alterations may depend on whether these alterations are secondary to increased network activity and thus reversible through homeostatic mechanisms [66].

## Declarations

### Ethics approval

Not applicable. Animals were killed solely to obtain tissue, which does not constitute a procedure under Directive 2010/63/EU.

### Consent to participate

Not applicable.

### Consent for publication

Not applicable.

### Data availability

The datasets generated and analyzed during the current study are available from the corresponding author on request.

### Code availability

Custom Python scripts used for data aggregation and selection of representative images are available from the corresponding author on request.

### Author contributions

MW conceived and designed the study, acquired funding, performed all experiments, analyzed most of the data, prepared figures, and wrote the manuscript. KK analyzed hippocampal dendritic spine data and prepared figures. PC performed Ca²⁺ imaging experiments together with MW. PM and KG analyzed part of the dendritic spine data. AA prepared primary cultures together with MW. AZ-M contributed to data interpretation. IN provided general supervision of the project. All authors read and approved the final manuscript.

## Acknowledgements

We thank Dr Bartłomiej Pochwat (Maj Institute of Pharmacology, Polish Academy of Sciences, Kraków, Poland) for providing ketamine.

## References

1. Nimchinsky EA, Sabatini BL, Svoboda K (2002) Structure and Function of Dendritic Spines. Annu Rev Physiol 64:313–353. 10.1146/annurev.physiol.64.081501.160008

2. Bourne JN, Harris KM (2008) Balancing Structure and Function at Hippocampal Dendritic Spines. Annu Rev Neurosci 31:47–67. 10.1146/annurev.neuro.31.060407.125646

3. Bonhoeffer T, Yuste R (2002) Spine Motility. Neuron 35:1019–1027. 10.1016/S0896-6273(02)00906-6

4. Matsuzaki M, Ellis-Davies GCR, Nemoto T, et al (2001) Dendritic spine geometry is critical for AMPA receptor expression in hippocampal CA1 pyramidal neurons. Nat Neurosci 4:1086–1092. 10.1038/nn736

5. Bourne J, Harris KM (2007) Do thin spines learn to be mushroom spines that remember? Curr Opin Neurobiol 17:381–386. 10.1016/j.conb.2007.04.009

6. Forrest MP, Parnell E, Penzes P (2018) Dendritic structural plasticity and neuropsychiatric disease. Nat Rev Neurosci 19:215–234. 10.1038/nrn.2018.16

7. McEwen BS, Nasca C, Gray JD (2016) Stress Effects on Neuronal Structure: Hippocampus, Amygdala, and Prefrontal Cortex. Neuropsychopharmacology 41:3–23. 10.1038/npp.2015.171

8. De Kloet ER, Vreugdenhil E, Oitzl MS, Joëls M (1998) Brain Corticosteroid Receptor Balance in Health and Disease. Endocr Rev 19:269–301. 10.1210/edrv.19.3.0331

9. Liston C, Cichon JM, Jeanneteau F, et al (2013) Circadian glucocorticoid oscillations promote learning-dependent synapse formation and maintenance. Nat Neurosci 16:698–705. 10.1038/nn.3387

10. Sarabdjitsingh RA, Jezequel J, Pasricha N, et al (2014) Ultradian corticosterone pulses balance glutamatergic transmission and synaptic plasticity. Proc Natl Acad Sci USA 111:14265–14270. 10.1073/pnas.1411216111

11. Gourley SL, Swanson AM, Koleske AJ (2013) Corticosteroid-Induced Neural Remodeling Predicts Behavioral Vulnerability and Resilience. J Neurosci 33:3107–3112. 10.1523/JNEUROSCI.2138-12.2013

12. Moda-Sava RN, Murdock MH, Parekh PK, et al (2019) Sustained rescue of prefrontal circuit dysfunction by antidepressant-induced spine formation. Science 364:eaat8078. 10.1126/science.aat8078

13. Tanokashira D, Morita T, Hayashi K, et al (2012) Glucocorticoid suppresses dendritic spine development mediated by down-regulation of caldesmon expression. J Neurosci 32:14583– 14591. 10.1523/JNEUROSCI.2380-12.2012

14. Radley JJ, Rocher AB, Rodriguez A, et al (2008) Repeated stress alters dendritic spine morphology in the rat medial prefrontal cortex. J Comp Neurol 507:1141–1150. 10.1002/cne.21588

15. Joëls M, Baram TZ (2009) The neuro-symphony of stress. Nat Rev Neurosci 10:459–466. 10.1038/nrn2632

16. Anderson RM, Glanz RM, Johnson SB, et al (2016) Prolonged corticosterone exposure induces dendritic spine remodeling and attrition in the rat medial prefrontal cortex. J Comp Neurol 524:3729–3746. 10.1002/cne.24027

17. Duman RS, Aghajanian GK (2012) Synaptic Dysfunction in Depression: Potential Therapeutic Targets. Science 338:68–72. 10.1126/science.1222939

18. Kang HJ, Voleti B, Hajszan T, et al (2012) Decreased expression of synapse-related genes and loss of synapses in major depressive disorder. Nat Med 18:1413–1417. 10.1038/nm.2886

19. Lamers F, Vogelzangs N, Merikangas KR, et al (2013) Evidence for a differential role of HPA-axis function, inflammation and metabolic syndrome in melancholic versus atypical depression. Mol Psychiatry 18:692–699. 10.1038/mp.2012.144

20. Holmes SE, Scheinost D, Finnema SJ, et al (2019) Lower synaptic density is associated with depression severity and network alterations. Nat Commun 10:1529. 10.1038/s41467-019-09562-7

21. Rush AJ, Trivedi MH, Wisniewski SR, et al (2006) Acute and Longer-Term Outcomes in Depressed Outpatients Requiring One or Several Treatment Steps: A STAR*D Report. Am J Psychiatry 163:1905–1917. 10.1176/ajp.2006.163.11.1905

22. Zarate CA, Singh JB, Carlson PJ, et al (2006) A Randomized Trial of an N-methyl-D-aspartate Antagonist in Treatment-Resistant Major Depression. Arch Gen Psychiatry 63:856–864. 10.1001/archpsyc.63.8.856

23. Murrough JW, Iosifescu DV, Chang LC, et al (2013) Antidepressant Efficacy of Ketamine in Treatment-Resistant Major Depression: A Two-Site Randomized Controlled Trial. Am J Psychiatry 170:1134–1142. 10.1176/appi.ajp.2013.13030392

24. Komatsuzaki Y, Hatanaka Y, Murakami G, et al (2012) Corticosterone induces rapid spinogenesis via synaptic glucocorticoid receptors and kinase networks in hippocampus. PLoS One 7:e34124. 10.1371/journal.pone.0034124

25. Ly C, Greb AC, Cameron LP, et al (2018) Psychedelics Promote Structural and Functional Neural Plasticity. Cell Rep 23:3170–3182. 10.1016/j.celrep.2018.05.022

26. Clark RD, Ray NC, Williams K, et al (2008) 1H-Pyrazolo[3,4-g]hexahydro-isoquinolines as selective glucocorticoid receptor antagonists with high functional activity. Bioorg Med Chem Lett 18:1312–1317. 10.1016/j.bmcl.2008.01.027

27. Zalachoras I, Houtman R, Atucha E, et al (2013) Differential targeting of brain stress circuits with a selective glucocorticoid receptor modulator. Proc Natl Acad Sci USA 110:7910–7915. 10.1073/pnas.1219411110

28. Du J, Wang Y, Hunter R, et al (2009) Dynamic regulation of mitochondrial function by glucocorticoids. Proc Natl Acad Sci USA 106:3543–3548. 10.1073/pnas.0812671106

29. Ruszczycki B, Szepesi Z, Wilczynski GM, et al (2012) Sampling issues in quantitative analysis of dendritic spines morphology. BMC Bioinformatics 13:213. 10.1186/1471-2105-13-213

30. Arshadi C, Günther U, Eddison M, et al (2021) SNT: a unifying toolbox for quantification of neuronal anatomy. Nat Methods 18:374–377. 10.1038/s41592-021-01105-7

31. Arganda-Carreras I, Kaynig V, Rueden C, et al (2017) Trainable Weka Segmentation: a machine learning tool for microscopy pixel classification. Bioinformatics 33:2424–2426. 10.1093/bioinformatics/btx180

32. Sutachan J-J, Montoya JV, Xu F, et al (2006) Pluronic F-127 affects the regulation of cytoplasmic Ca2+ in neuronal cells. Brain Res 1068:131–137. 10.1016/j.brainres.2005.11.003

33. Patel TP, Man K, Firestein BL, Meaney DF (2015) Automated quantification of neuronal networks and single-cell calcium dynamics using calcium imaging. J Neurosci Methods 243:26–38. 10.1016/j.jneumeth.2015.01.020

34. Stringer C, Wang T, Michaelos M, Pachitariu M (2021) Cellpose: a generalist algorithm for cellular segmentation. Nat Methods 18:100–106. 10.1038/s41592-020-01018-x

35. Livak KJ, Schmittgen TD (2001) Analysis of Relative Gene Expression Data Using Real-Time Quantitative PCR and the 2−ΔΔCT Method. Methods 25:402–408. 10.1006/meth.2001.1262

36. Digilio L, Yap CC, Winckler B (2015) Ctip2-, Satb2-, Prox1-, and GAD65-Expressing Neurons in Rat Cultures: Preponderance of Single-and Double-Positive Cells, and Cell Type-Specific Expression of Neuron-Specific Gene Family Members, Nsg-1 (NEEP21) and Nsg-2 (P19). PLoS One 10:e0140010. 10.1371/journal.pone.0140010

37. Wyrwoll CS, Holmes MC, Seckl JR (2011) 11β-Hydroxysteroid dehydrogenases and the brain: From zero to hero, a decade of progress. Front Neuroendocrinol 32:265–286. 10.1016/j.yfrne.2010.12.001

38. Bassil K, Krontira AC, Leroy T, et al (2023) In vitro modeling of the neurobiological effects of glucocorticoids: A review. Neurobiol Stress 23:100530. 10.1016/j.ynstr.2023.100530

39. Holtmaat AJGD, Trachtenberg JT, Wilbrecht L, et al (2005) Transient and persistent dendritic spines in the neocortex in vivo. Neuron 45:279–291. 10.1016/j.neuron.2005.01.003

40. Meyer D, Bonhoeffer T, Scheuss V (2014) Balance and Stability of Synaptic Structures during Synaptic Plasticity. Neuron 82:430–443. 10.1016/j.neuron.2014.02.031

41. Chen X, Vinade L, Leapman RD, et al (2005) Mass of the postsynaptic density and enumeration of three key molecules. Proc Natl Acad Sci USA 102:11551–11556. 10.1073/pnas.0505359102

42. Pchitskaya E, Bezprozvanny I (2020) Dendritic Spines Shape Analysis—Classification or Clusterization? Perspective. Front Synaptic Neurosci 12:31. 10.3389/fnsyn.2020.00031

43. Koo JW, Wohleb ES (2021) How Stress Shapes Neuroimmune Function: Implications for the Neurobiology of Psychiatric Disorders. Biol Psychiatry 90:74–84. 10.1016/j.biopsych.2020.11.007

44. Wohleb ES, Terwilliger R, Duman CH, Duman RS (2018) Stress-Induced Neuronal Colony Stimulating Factor 1 Provokes Microglia-Mediated Neuronal Remodeling and Depressive-like Behavior. Biol Psychiatry 83:38–49. 10.1016/j.biopsych.2017.05.026

45. Tillmon H, Soteros BM, Shen L, et al (2024) Complement and microglia activation mediate stress-induced synapse loss in layer 2/3 of the medial prefrontal cortex in male mice. Nat Commun 15:9803. 10.1038/s41467-024-54007-5

46. Horchar MJ, Wohleb ES (2019) Glucocorticoid receptor antagonism prevents microglia-mediated neuronal remodeling and behavioral despair following chronic unpredictable stress. Brain Behav Immun 81:329–340. 10.1016/j.bbi.2019.06.030

47. Radley JJ, Anderson RM, Hamilton BA, et al (2013) Chronic Stress-Induced Alterations of Dendritic Spine Subtypes Predict Functional Decrements in an Hypothalamo–Pituitary–Adrenal-Inhibitory Prefrontal Circuit. J Neurosci 33:14379–14391. 10.1523/JNEUROSCI.0287-13.2013

48. Harris KM, Stevens JK (1989) Dendritic spines of CA 1 pyramidal cells in the rat hippocampus: serial electron microscopy with reference to their biophysical characteristics. J Neurosci 9:2982–2997. 10.1523/JNEUROSCI.09-08-02982.1989

49. Arellano JI (2007) Ultrastructure of dendritic spines: correlation between synaptic and spine morphologies. Front Neurosci 1:131–143. 10.3389/neuro.01.1.1.010.2007

50. Ehrlich I, Klein M, Rumpel S, Malinow R (2007) PSD-95 is required for activity-driven synapse stabilization. Proc Natl Acad Sci USA 104:4176–4181. 10.1073/pnas.0609307104

51. Cane M, Maco B, Knott G, Holtmaat A (2014) The Relationship between PSD-95 Clustering and Spine Stability *In Vivo*. J Neurosci 34:2075–2086. 10.1523/JNEUROSCI.3353-13.2014

52. Boyer C, Schikorski T, Stevens CF (1998) Comparison of Hippocampal Dendritic Spines in Culture and in Brain. J Neurosci 18:5294–5300. 10.1523/JNEUROSCI.18-14-05294.1998

53. Prange O, Murphy TH (2001) Modular Transport of Postsynaptic Density-95 Clusters and Association with Stable Spine Precursors during Early Development of Cortical Neurons. J Neurosci 21:9325–9333. 10.1523/JNEUROSCI.21-23-09325.2001

54. Woods GF, Oh WC, Boudewyn LC, et al (2011) Loss of PSD-95 Enrichment Is Not a Prerequisite for Spine Retraction. J Neurosci 31:12129–12138. 10.1523/JNEUROSCI.6662-10.2011

55. Fukata Y, Dimitrov A, Boncompain G, et al (2013) Local palmitoylation cycles define activity-regulated postsynaptic subdomains. J Cell Biol 202:145–161. 10.1083/jcb.201302071

56. Pytyś A, Ijaz R, Buszka A, et al (2025) Temporal and protein-specific S-palmitoylation supports synaptic and neural network plasticity. Cell Mol Life Sci 82:355. 10.1007/s00018-025-05893-5

57. Yuen EY, Liu W, Karatsoreos IN, et al (2011) Mechanisms for acute stress-induced enhancement of glutamatergic transmission and working memory. Mol Psychiatry 16:156–170. 10.1038/mp.2010.50

58. Yuen EY, Wei J, Liu W, et al (2012) Repeated Stress Causes Cognitive Impairment by Suppressing Glutamate Receptor Expression and Function in Prefrontal Cortex. Neuron 73:962–977. 10.1016/j.neuron.2011.12.033

59. Kirov SA, Harris KM (1999) Dendrites are more spiny on mature hippocampal neurons when synapses are inactivated. Nat Neurosci 2:878–883. 10.1038/13178

60. Wefelmeyer W, Puhl CJ, Burrone J (2016) Homeostatic Plasticity of Subcellular Neuronal Structures: From Inputs to Outputs. Trends Neurosci 39:656–667. 10.1016/j.tins.2016.08.004

61. Karst H, Berger S, Turiault M, et al (2005) Mineralocorticoid receptors are indispensable for nongenomic modulation of hippocampal glutamate transmission by corticosterone. Proc Natl Acad Sci USA 102:19204–19207. 10.1073/pnas.0507572102

62. Turrigiano GG, Leslie KR, Desai NS, et al (1998) Activity-dependent scaling of quantal amplitude in neocortical neurons. Nature 391:892–896. 10.1038/36103

63. Lee KJ, Lee Y, Rozeboom A, et al (2011) Requirement for Plk2 in Orchestrated Ras and Rap Signaling, Homeostatic Structural Plasticity, and Memory. Neuron 69:957–973. 10.1016/j.neuron.2011.02.004

64. Reul JMHM, de Kloet ER (1985) Two Receptor Systems for Corticosterone in Rat Brain: Microdistribution and Differential Occupation. Endocrinology 117:2505–2511. 10.1210/endo-117-6-2505

65. Autry AE, Adachi M, Nosyreva E, et al (2011) NMDA receptor blockade at rest triggers rapid behavioural antidepressant responses. Nature 475:91–95. 10.1038/nature10130

66. Kavalali ET, Monteggia LM (2020) Targeting Homeostatic Synaptic Plasticity for Treatment of Mood Disorders. Neuron 106:715–726. 10.1016/j.neuron.2020.05.015

67. Ng LHL, Huang Y, Han L, et al (2018) Ketamine and selective activation of parvalbumin interneurons inhibit stress-induced dendritic spine elimination. Transl Psychiatry 8:272. 10.1038/s41398-018-0321-5

68. Teppola H, Aćimović J, Linne M-L (2019) Unique Features of Network Bursts Emerge From the Complex Interplay of Excitatory and Inhibitory Receptors in Rat Neocortical Networks. Front Cell Neurosci 13:377. 10.3389/fncel.2019.00377

69. Ahtiainen A, Annala I, Rosenholm M, et al (2023) Ketamine reduces electrophysiological network activity in cortical neuron cultures already at sub-micromolar concentrations – Impact on TrkB-ERK1/2 signaling. Neuropharmacology 229:109481. 10.1016/j.neuropharm.2023.109481

70. Masuda FK, Aery Jones EA, Sun Y, Giocomo LM (2023) Ketamine evoked disruption of entorhinal and hippocampal spatial maps. Nat Commun 14:6285. 10.1038/s41467-023-41750-4

71. Cichon J, Wasilczuk AZ, Looger LL, et al (2023) Ketamine triggers a switch in excitatory neuronal activity across neocortex. Nat Neurosci 26:39–52. 10.1038/s41593-022-01203-5

