## Supplementary Material for "Ketamine reverses chronic corticosterone-induced changes in dendritic spine morphology in hippocampal but not cortical neurons in vitro"

**Contents**

Supplementary Methods
Supplementary Fig. S1 – Cellular composition and regional identity of primary cultures
Supplementary Fig. S2 – Additional parameters of spontaneous $Ca^{2+}$ activity
Supplementary Table S1 – Summary of statistical analyses

### Supplementary Methods

#### Coverslip and plate preparation

Primary cortical and hippocampal cultures were grown on 18-mm glass coverslips (Fisher Scientific, cat. no. 11817093), clear-bottom 96-well imaging plates (Revvity, cat. no. 605530), or 6-well plates (Bionovo, cat. no. B-0680), depending on the experiment. Coverslips, used only to evaluate the effects of different CORT concentrations on spine morphology, were prepared as follows. First, they were immersed in 70% nitric acid for 24 h. Then, they were washed seven times in deionized water (90 min in total) and sonicated in fresh deionized water for 30 min. Finally, the coverslips were sterilized by autoclaving followed by a rinse in anhydrous ethanol (POL-AURA, cat. no. PA-16-6480111), air-dried, and coated overnight at 4 °C with poly-D-lysine (PDL; 0.5 mg/ml; Sigma-Aldrich, cat. no. P0899) diluted in PBS (Sigma-Aldrich, cat. no. P4417). The next day, they were washed three times with PBS, air-dried, transferred to 12-well plates (Bionovo, cat. no. B-0681), and stored in a laminar flow hood until use. The wells of the 96- and 6-well plates were coated with PDL (0.1 mg/ml) as above. The next day, the PDL solution was removed, and the wells were washed three times with PBS and air-dried. The plates were then stored in a laminar flow hood until use.

#### Quantification of the proportions of NeuN$^+$ and S100B$^+$ cells

Images for quantifying the proportions of NeuN$^+$ and S100B$^+$ cells were acquired at 2048 × 2048 pixels with 470 nm excitation and a 300 ms exposure time for both markers, which were stained in separate, parallel wells. Images used for cell counting were processed with Large Volume Computational Clearing (LVCC; Leica) using identical settings for all images, and the processed images were analyzed. To quantify the proportions of NeuN$^+$ and S100B$^+$ cells, NeuN$^+$ cells, S100B$^+$ cells, and Hoechst-stained nuclei were counted using a batch macro based on the Trainable Weka Segmentation plugin [Arganda-Carreras et al. 2017]. Each image was converted to 8-bit and background-subtracted (rolling ball, radius 50 pixels). Separate classifiers, trained on images preprocessed in the same way, were used for NeuN, S100B, and Hoechst to distinguish intact cells from condensed, uniformly bright (pyknotic) nuclei and debris and to generate class probability maps. The probability map of condensed nuclei and debris was subtracted from that of intact cells, and the resulting map was thresholded using the Li method, median-filtered (radius 4 pixels), and segmented with the watershed algorithm. Objects ≥ 30 µm² with a circularity of 0.30–1.00 were counted, excluding objects touching the image border. For each marker, the same classifier and parameters were applied to all images. The proportions of NeuN$^+$ and S100B$^+$ cells were calculated as the number of NeuN$^+$ or S100B$^+$ cells divided by the number of intact nuclei in the same image; NeuN and S100B were assessed in separate, parallel wells. The proportions of NeuN$^+$ and S100B$^+$ cells were determined in one well per marker (25 fields per well) from one culture.

#### References

Arganda-Carreras I, Kaynig V, Rueden C, Eliceiri KW, Schindelin J, Cardona A, Seung HS (2017) Trainable Weka Segmentation: a machine learning tool for microscopy pixel classification. Bioinformatics 33:2424–2426. https://doi.org/10.1093/bioinformatics/btx180

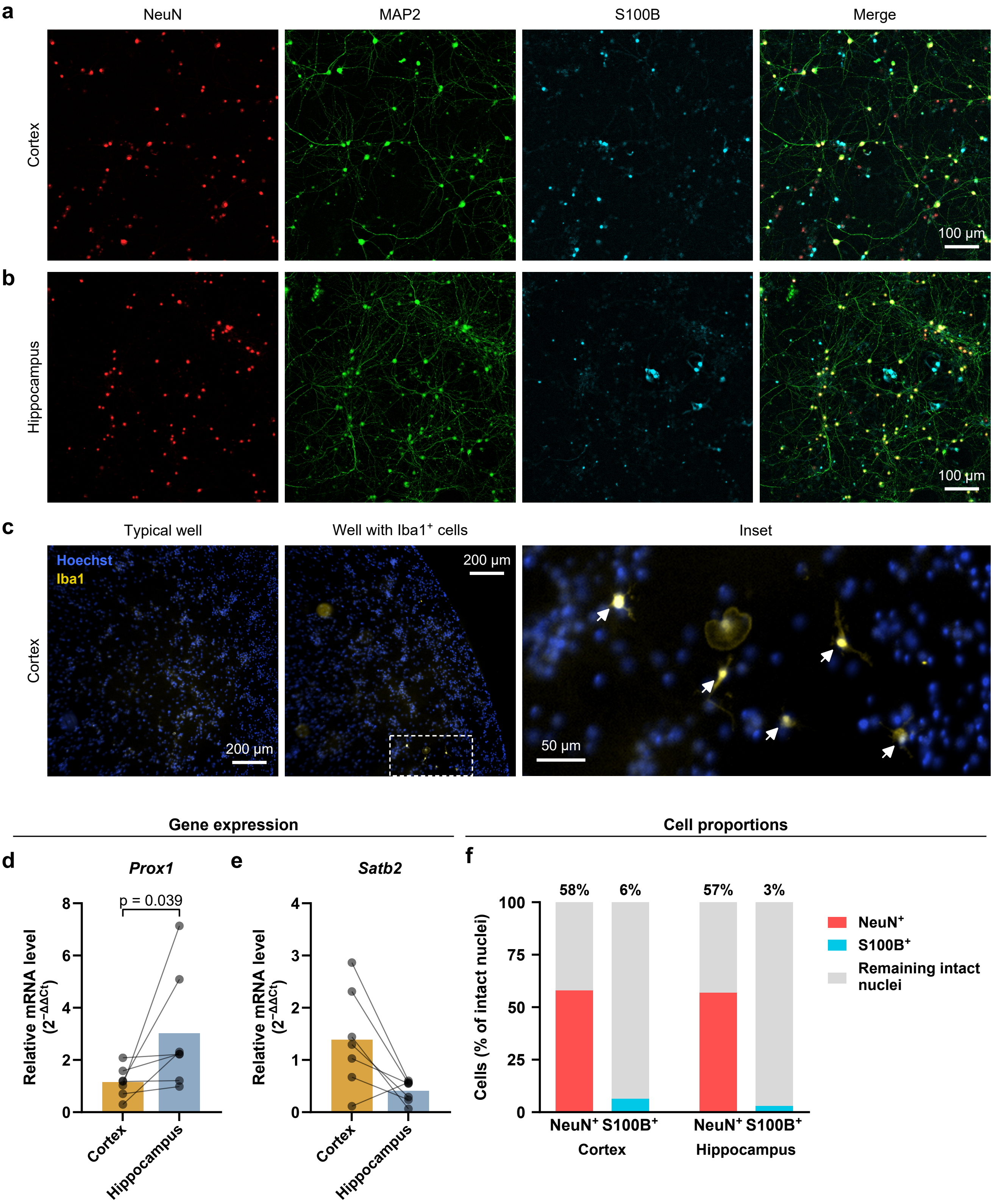
a
NeuN
MAP2
S100B
Merge
Cortex
100 µm
b
Hippocampus
100 µm
c
Typical well
Well with Iba1+ cells
Inset
Hoechst
Iba1
Cortex
200 µm
200 µm
50 µm
Gene expression
Cell proportions
d
Prox1
p = 0.039
Relative mRNA level (2^−ΔΔCt)
Cortex
Hippocampus
e
Satb2
f
58%
6%
57%
3%
Cells (% of intact nuclei)
NeuN+ S100B+
Cortex
Hippocampus
NeuN+
S100B+
Remaining intact nuclei

**Supplementary Fig. S1** Cellular composition and regional identity of primary cultures. **a, b** Representative images (not used for quantifications) of cortical (**a**) and hippocampal (**b**) cultures at DIV18 immunostained for NeuN (red), MAP2 (green), and S100B (cyan). **c** Iba1 (yellow) and Hoechst 33342 (blue) staining in cortical cultures: a typical well, a well containing $Iba1^+$ cells, and an enlargement of the boxed area (arrows, $Iba1^+$ cells). **d, e** *Prox1* (**d**) and *Satb2* (**e**) mRNA levels in hippocampal relative to cortical cultures (n = 7 cultures). In **d** and **e**, bars represent arithmetic means of $2^{-\Delta\Delta Ct}$ values, whereas statistical analyses were performed on log-transformed values and geometric mean fold changes are given in the text. **f** Proportions of $NeuN^+$ and $S100B^+$ cells among intact nuclei (one well per marker, 25 fields per well, one culture). Points represent individual cultures, and lines connect values from the same culture. The p value from the ratio paired t-test is shown for the significant comparison. Scale bars: 100 µm (**a, b**), 200 µm (**c**, whole wells), 50 µm (**c**, enlargement)

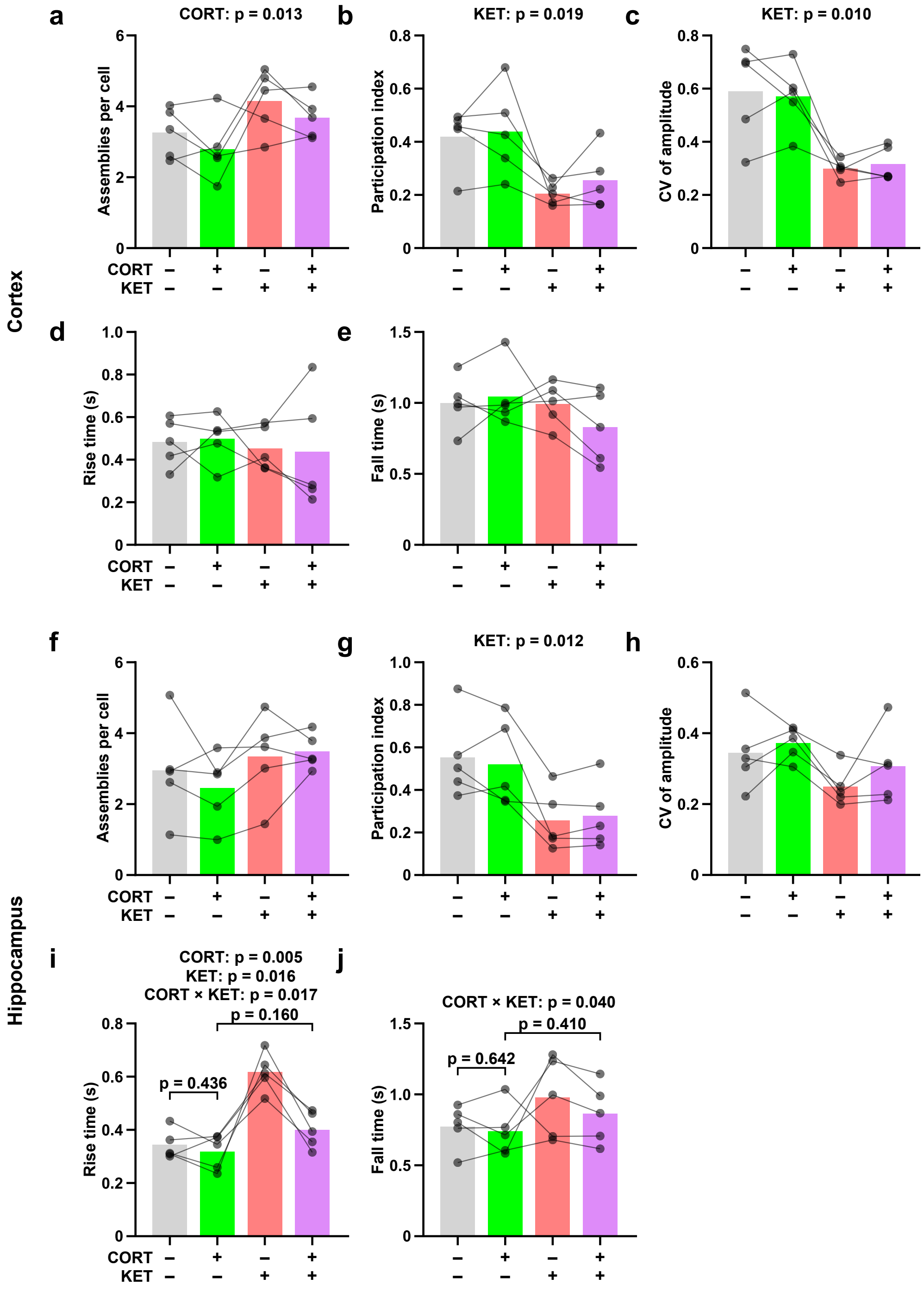


**Supplementary Fig. S2** Additional parameters of spontaneous $Ca^{2+}$ activity in cortical (**a–e**) and hippocampal (**f–j**) cultures. Number of assemblies per cell (**a**, **f**), participation index (**b**, **g**), coefficient of variation of transient amplitude (**c**, **h**), rise time (**d**, **i**), and fall time (**e**, **j**) of $Ca^{2+}$ transients; n = 5 cultures. Bars represent means, points represent individual cultures, and lines connect values from the same culture. Significant main effects and interactions from two-way RM ANOVA are shown above the graphs; for a significant interaction, p values from planned paired t-tests (DMSO vs CORT; CORT vs CORT + KET) are shown

**Supplementary Table S1** Summary of statistical analyses. Test statistics, degrees of freedom, and p values for all analyses reported in the Results, listed by figure panel (Fig. 1–7, Supplementary Fig. S1, S2). Main effects and interactions are from one-, two-, or three-way RM (mixed) ANOVA or mixed-effects models, as indicated; post hoc (Dunnett vs DMSO) and planned or exploratory paired t-tests are listed below the corresponding main analysis. n is given in the figure legends.

| Panel | Culture | Parameter | Analysis | Effect / comparison | Statistic | p |
|---|---|---|---|---|---|---|
| 1c | Cortex | *Fkbp5* mRNA | Mixed-effects model | CORT | F(3, 10) = 42.88 | < 0.0001 |
| | | | | Dunnett: 0.25 µM vs DMSO | | < 0.0001 |
| | | | | Dunnett: 0.5 µM vs DMSO | | < 0.0001 |
| | | | | Dunnett: 1 µM vs DMSO | | < 0.0001 |
| 1d | Hippocampus | *Fkbp5* mRNA | Mixed-effects model | CORT | F(3, 12) = 3.77 | 0.040 |
| | | | | Dunnett: 0.25 µM vs DMSO | | 0.213 |
| | | | | Dunnett: 0.5 µM vs DMSO | | 0.135 |
| | | | | Dunnett: 1 µM vs DMSO | | 0.018 |
| 1e | Cortex | LDH release | One-way RM ANOVA | CORT | F(3, 9) = 2.29 | 0.146 |
| 1f | Hippocampus | LDH release | One-way RM ANOVA | CORT | F(3, 9) = 6.40 | 0.013 |
| | | | | Dunnett: 0.25 µM vs DMSO | | 0.211 |
| | | | | Dunnett: 0.5 µM vs DMSO | | 0.819 |
| | | | | Dunnett: 1 µM vs DMSO | | 0.007 |
| 1g | Cortex | NeuN$^{+}$ cells | One-way RM ANOVA | CORT | F(3, 6) = 0.99 | 0.459 |
| 1h | Hippocampus | NeuN$^{+}$ cells | One-way RM ANOVA | CORT | F(3, 6) = 2.46 | 0.160 |
| 1j | Cortex | Total dendritic length | One-way RM ANOVA | CORT | F(3, 6) = 41.40 | 0.0002 |
| | | | | Dunnett: 0.25 µM vs DMSO | | 0.379 |
| | | | | Dunnett: 0.5 µM vs DMSO | | 0.0007 |
| | | | | Dunnett: 1 µM vs DMSO | | 0.0002 |
| 1k | Hippocampus | Total dendritic length | One-way RM ANOVA | CORT | F(3, 6) = 10.11 | 0.009 |
| | | | | Dunnett: 0.25 µM vs DMSO | | 0.359 |
| | | | | Dunnett: 0.5 µM vs DMSO | | 0.042 |
| | | | | Dunnett: 1 µM vs DMSO | | 0.004 |
| Fig. S1d | Hippocampus vs cortex | *Prox1* mRNA | Ratio paired t-test | Region | t(6) = 2.62 | 0.039 |
| Fig. S1e | Hippocampus vs cortex | *Satb2* mRNA | Ratio paired t-test | Region | t(6) = 2.05 | 0.086 |
| 2b | Cortex | Spine density | One-way RM ANOVA | CORT | F(3, 6) = 3.33 | 0.097 |
| 2c | Cortex | Spine length | One-way RM ANOVA | CORT | F(3, 6) = 3.11 | 0.109 |
| 2d | Cortex | Spine head width | One-way RM ANOVA | CORT | F(3, 6) = 23.0 | 0.001 |
| | | | | Dunnett: 0.25 µM vs DMSO | | 0.004 |
| | | | | Dunnett: 0.5 µM vs DMSO | | 0.0008 |
| | | | | Dunnett: 1 µM vs DMSO | | 0.001 |
| 2e | Cortex | Length-to-head-width ratio | One-way RM ANOVA | CORT | F(3, 6) = 24.5 | 0.0009 |
| | | | | Dunnett: 0.25 µM vs DMSO | | 0.002 |
| | | | | Dunnett: 0.5 µM vs DMSO | | 0.001 |
| | | | | Dunnett: 1 µM vs DMSO | | 0.0005 |

**Supplementary Table S1** (continued)

| Panel | Culture | Parameter | Analysis | Effect / comparison | Statistic | p |
|---|---|---|---|---|---|---|
| 2f | Cortex | Spine area | One-way RM ANOVA | CORT | $F(3, 6) = 5.01$ | 0.044 |
| | | | | Dunnett: 0.25 µM vs DMSO | | 0.436 |
| | | | | Dunnett: 0.5 µM vs DMSO | | 0.025 |
| | | | | Dunnett: 1 µM vs DMSO | | 0.093 |
| 2h | Hippocampus | Spine density | One-way RM ANOVA | CORT | $F(3, 6) = 5.68$ | 0.034 |
| | | | | Dunnett: 0.25 µM vs DMSO | | 0.048 |
| | | | | Dunnett: 0.5 µM vs DMSO | | 0.024 |
| | | | | Dunnett: 1 µM vs DMSO | | 0.047 |
| 2i | Hippocampus | Spine length | One-way RM ANOVA | CORT | $F(3, 6) = 11.86$ | 0.006 |
| | | | | Dunnett: 0.25 µM vs DMSO | | 0.005 |
| | | | | Dunnett: 0.5 µM vs DMSO | | 0.005 |
| | | | | Dunnett: 1 µM vs DMSO | | 0.022 |
| 2j | Hippocampus | Spine head width | One-way RM ANOVA | CORT | $F(3, 6) = 21.48$ | 0.001 |
| | | | | Dunnett: 0.25 µM vs DMSO | | 0.001 |
| | | | | Dunnett: 0.5 µM vs DMSO | | 0.005 |
| | | | | Dunnett: 1 µM vs DMSO | | 0.0009 |
| 2k | Hippocampus | Length-to-head-width ratio | One-way RM ANOVA | CORT | $F(3, 6) = 2.33$ | 0.173 |
| 2l | Hippocampus | Spine area | One-way RM ANOVA | CORT | $F(3, 6) = 33.0$ | 0.0004 |
| | | | | Dunnett: 0.25 µM vs DMSO | | 0.0004 |
| | | | | Dunnett: 0.5 µM vs DMSO | | 0.0007 |
| | | | | Dunnett: 1 µM vs DMSO | | 0.0004 |
| 2n | Cortex (culture medium) | Spine density | Mixed-effects model / Planned paired t-test | Culture condition | $F(2, 8) = 25.47$ | 0.0003 |
| | | | | MEM/N2 vs NB/B27 | $t(3) = 4.29$ | 0.023 |
| | | | | Paired t-test: NB/B27 vs NB/B27 + CORT | $t(2) = 1.59$ | 0.253 |
| 2o | Cortex (culture medium) | Spine length | Mixed-effects model / Planned paired t-test | Culture condition | $F(2, 8) = 18.71$ | 0.001 |
| | | | | MEM/N2 vs NB/B27 | $t(3) = 3.21$ | 0.049 |
| | | | | Paired t-test: NB/B27 vs NB/B27 + CORT | $t(2) = 2.24$ | 0.155 |
| 2p | Cortex (culture medium) | Spine head width | Mixed-effects model / Planned paired t-test | Culture condition | $F(2, 5) = 29.33$ | 0.002 |
| | | | | MEM/N2 vs NB/B27 | $t(3) = 5.10$ | 0.015 |
| | | | | Paired t-test: NB/B27 vs NB/B27 + CORT | $t(2) = 1.08$ | 0.393 |

**Supplementary Table S1** (continued)

| Panel | Culture | Parameter | Analysis | Effect / comparison | Statistic | p |
|---|---|---|---|---|---|---|
| 2q | Cortex (culture medium) | Length-to-head-width ratio | Mixed-effects model / Planned paired t-test | Culture condition | F(2, 5) = 87.53 | 0.0001 |
| | | | | MEM/N2 vs NB/B27 | t(3) = 9.64 | 0.002 |
| | | | | Paired t-test: NB/B27 vs NB/B27 + CORT | t(2) = 3.63 | 0.068 |
| 2r | Cortex (culture medium) | Spine area | Mixed-effects model | Culture condition | F(2, 5) = 0.540 | 0.613 |
| 3b | Cortex | PSD-95 puncta density | Paired t-test | DMSO vs CORT | t(2) = 1.24 | 0.340 |
| 3c | Cortex | PSD-95 puncta size | Paired t-test | DMSO vs CORT | t(2) = 0.70 | 0.555 |
| 3d | Cortex | Mean PSD-95 intensity | Paired t-test | DMSO vs CORT | t(2) = 1.19 | 0.353 |
| 3f | Hippocampus | PSD-95 puncta density | Paired t-test | DMSO vs CORT | t(2) = 1.53 | 0.263 |
| 3g | Hippocampus | PSD-95 puncta size | Paired t-test | DMSO vs CORT | t(2) = 0.20 | 0.859 |
| 3h | Hippocampus | Mean PSD-95 intensity | Paired t-test | DMSO vs CORT | t(2) = 0.300 | 0.792 |
| 4b | Cortex | *Fkbp5* mRNA | Two-way RM ANOVA | CORT | F(1, 4) = 92.27 | 0.0007 |
| | | | | C108 | F(1, 4) = 0.54 | 0.504 |
| | | | | CORT × C108 | F(1, 4) = 5.17 | 0.085 |
| 4c | Cortex | Spine density | Two-way RM ANOVA | CORT | F(1, 3) = 1.00 | 0.391 |
| | | | | C108 | F(1, 3) = 4.44 | 0.126 |
| | | | | CORT × C108 | F(1, 3) = 0.58 | 0.501 |
| 4d | Cortex | Spine length | Two-way RM ANOVA | CORT | F(1, 3) = 1.42 | 0.319 |
| | | | | C108 | F(1, 3) = 1.14 | 0.363 |
| | | | | CORT × C108 | F(1, 3) = 0.041 | 0.853 |
| 4e | Cortex | Spine head width | Two-way RM ANOVA | CORT × C108 | F(1, 3) = 12.94 | 0.037 |
| | | | | CORT | F(1, 3) = 10.05 | 0.051 |
| | | | | C108 | F(1, 3) = 1.48 | 0.310 |
| | | | | Planned t-test: DMSO vs CORT | t(3) = 4.03 | 0.028 |
| | | | | Planned t-test: CORT vs CORT + C108 | t(3) = 4.17 | 0.025 |
| | | | | Exploratory t-test: C108 vs CORT + C108 | t(3) = 0.18 | 0.866 |
| | | | | Exploratory t-test: DMSO vs C108 | t(3) = 2.55 | 0.084 |
| 4f | Cortex | Length-to-head-width ratio | Two-way RM ANOVA | CORT | F(1, 3) = 15.23 | 0.030 |
| | | | | C108 | F(1, 3) = 1.16 | 0.361 |
| | | | | CORT × C108 | F(1, 3) = 1.62 | 0.293 |
| 4g | Cortex | Spine area | Two-way RM ANOVA | CORT | F(1, 3) = 0.16 | 0.718 |
| | | | | C108 | F(1, 3) = 1.29 | 0.338 |
| | | | | CORT × C108 | F(1, 3) = 4.57 | 0.122 |
| 4i | Hippocampus | Spine density | Two-way RM ANOVA | CORT | F(1, 3) = 3.62 | 0.153 |
| | | | | C108 | F(1, 3) = 1.11 | 0.370 |
| | | | | CORT × C108 | F(1, 3) = 1.37 | 0.326 |

**Supplementary Table S1** (continued)

| Panel | Culture | Parameter | Analysis | Effect / comparison | Statistic | p |
|---|---|---|---|---|---|---|
| 4j | Hippocampus | Spine length | Two-way RM ANOVA | CORT | F(1, 3) = 14.37 | 0.032 |
| | | | | C108 | F(1, 3) = 0.13 | 0.747 |
| | | | | CORT × C108 | F(1, 3) = 3.82 | 0.146 |
| 4k | Hippocampus | Spine head width | Two-way RM ANOVA | CORT | F(1, 3) = 36.69 | 0.009 |
| | | | | C108 | F(1, 3) = 0.002 | 0.968 |
| | | | | CORT × C108 | F(1, 3) = 5.21 | 0.107 |
| 4l | Hippocampus | Length-to-head-width ratio | Two-way RM ANOVA | CORT | F(1, 3) = 7.00 | 0.077 |
| | | | | C108 | F(1, 3) = 0.21 | 0.678 |
| | | | | CORT × C108 | F(1, 3) = 2.07 | 0.246 |
| 4m | Hippocampus | Spine area | Two-way RM ANOVA | CORT | F(1, 3) = 23.40 | 0.017 |
| | | | | C108 | F(1, 3) = 0.02 | 0.888 |
| | | | | CORT × C108 | F(1, 3) = 9.22 | 0.056 |
| 4 | Cortex vs hippocampus | Spine density | Three-way mixed ANOVA | Region × CORT × C108 | F(1, 6) = 0.51 | 0.502 |
| 4 | Cortex vs hippocampus | Spine length | Three-way mixed ANOVA | Region × CORT × C108 | F(1, 6) = 1.47 | 0.271 |
| 4 | Cortex vs hippocampus | Spine head width | Three-way mixed ANOVA | Region × CORT × C108 | F(1, 6) = 0.06 | 0.820 |
| 4 | Cortex vs hippocampus | Length-to-head-width ratio | Three-way mixed ANOVA | Region × CORT × C108 | F(1, 6) = 2.44 | 0.169 |
| 4 | Cortex vs hippocampus | Spine area | Three-way mixed ANOVA | Region × CORT × C108 | F(1, 6) = 0.92 | 0.374 |
| 5b | Cortex | Spine density | Two-way RM ANOVA | CORT | F(1, 4) = 3.48 | 0.135 |
| | | | | KET | F(1, 4) = 0.018 | 0.901 |
| | | | | CORT × KET | F(1, 4) = 0.74 | 0.439 |
| 5c | Cortex | Spine length | Two-way RM ANOVA | KET | F(1, 4) = 12.60 | 0.024 |
| | | | | CORT | F(1, 4) < 0.001 | > 0.99 |
| | | | | CORT × KET | F(1, 4) = 7.27 | 0.054 |
| 5d | Cortex | Spine head width | Two-way RM ANOVA | CORT | F(1, 4) = 8.75 | 0.042 |
| | | | | KET | F(1, 4) = 1.71 | 0.261 |
| | | | | CORT × KET | F(1, 4) = 3.41 | 0.139 |
| 5e | Cortex | Length-to-head-width ratio | Two-way RM ANOVA | CORT × KET | F(1, 4) = 15.13 | 0.018 |
| | | | | CORT | F(1, 4) = 4.70 | 0.096 |
| | | | | KET | F(1, 4) = 1.23 | 0.330 |
| | | | | Planned t-test: DMSO vs CORT | t(4) = 4.32 | 0.012 |
| | | | | Planned t-test: CORT vs CORT + KET | t(4) = 2.06 | 0.109 |
| | | | | Exploratory t-test: DMSO vs KET | t(4) = 4.66 | 0.010 |
| 5f | Cortex | Spine area | Two-way RM ANOVA | CORT | F(1, 4) = 5.03 | 0.088 |
| | | | | KET | F(1, 4) = 3.80 | 0.123 |
| | | | | CORT × KET | F(1, 4) = 3.32 | 0.143 |
| 5h | Hippocampus | Spine density | Two-way RM ANOVA | CORT | F(1, 4) = 0.007 | 0.938 |
| | | | | KET | F(1, 4) = 0.68 | 0.457 |
| | | | | CORT × KET | F(1, 4) = 1.00 | 0.374 |

**Supplementary Table S1** (continued)

| Panel | Culture | Parameter | Analysis | Effect / comparison | Statistic | p |
|---|---|---|---|---|---|---|
| 5i | Hippocampus | Spine length | Two-way RM ANOVA | CORT × KET | F(1, 4) = 8.07 | 0.047 |
| | | | | CORT | F(1, 4) = 4.27 | 0.108 |
| | | | | KET | F(1, 4) = 5.57 | 0.078 |
| | | | | Planned t-test: DMSO vs CORT | t(4) = 3.43 | 0.027 |
| | | | | Planned t-test: CORT vs CORT + KET | t(4) = 6.09 | 0.004 |
| | | | | Exploratory t-test: DMSO vs CORT + KET | t(4) = 0.85 | 0.441 |
| | | | | Exploratory t-test: DMSO vs KET | t(4) = 0.22 | 0.833 |
| 5j | Hippocampus | Spine head width | Two-way RM ANOVA | CORT × KET | F(1, 4) = 38.81 | 0.003 |
| | | | | CORT | F(1, 4) = 2.13 | 0.218 |
| | | | | KET | F(1, 4) = 32.62 | 0.005 |
| | | | | Planned t-test: DMSO vs CORT | t(4) = 4.03 | 0.016 |
| | | | | Planned t-test: CORT vs CORT + KET | t(4) = 12.18 | 0.0003 |
| | | | | Exploratory t-test: DMSO vs CORT + KET | t(4) = 3.38 | 0.028 |
| | | | | Exploratory t-test: DMSO vs KET | t(4) = 1.31 | 0.261 |
| 5k | Hippocampus | Length-to-head-width ratio | Two-way RM ANOVA | CORT | F(1, 4) = 2.88 | 0.165 |
| | | | | KET | F(1, 4) = 1.23 | 0.329 |
| | | | | CORT × KET | F(1, 4) = 0.031 | 0.869 |
| 5l | Hippocampus | Spine area | Two-way RM ANOVA | CORT × KET | F(1, 4) = 38.28 | 0.003 |
| | | | | CORT | F(1, 4) = 2.99 | 0.159 |
| | | | | KET | F(1, 4) = 100.56 | 0.0006 |
| | | | | Planned t-test: DMSO vs CORT | t(4) = 3.77 | 0.020 |
| | | | | Planned t-test: CORT vs CORT + KET | t(4) = 19.69 | < 0.0001 |
| | | | | Exploratory t-test: DMSO vs CORT + KET | t(4) = 0.72 | 0.513 |
| | | | | Exploratory t-test: DMSO vs KET | t(4) = 1.16 | 0.311 |
| 5 | Cortex vs hippocampus | Spine head width | Three-way mixed ANOVA | Region × CORT × KET | F(1, 8) = 6.88 | 0.031 |
| 5 | Cortex vs hippocampus | Spine length | Three-way mixed ANOVA | Region × CORT × KET | F(1, 8) = 15.28 | 0.005 |
| 5 | Cortex vs hippocampus | Spine area | Three-way mixed ANOVA | Region × CORT × KET | F(1, 8) = 33.78 | < 0.001 |
| 5 | Cortex vs hippocampus | Length-to-head-width ratio | Three-way mixed ANOVA | Region × CORT × KET | F(1, 8) = 8.19 | 0.021 |
| 6d | Cortex | ROIs per field | Two-way RM ANOVA | CORT | F(1, 4) = 33.70 | 0.004 |
| | | | | KET | F(1, 4) = 14.04 | 0.020 |
| | | | | CORT × KET | F(1, 4) = 0.14 | 0.729 |
| 6e | Cortex | Active ROIs (%) | Two-way RM ANOVA | KET | F(1, 4) = 7.27 | 0.054 |
| | | | | CORT | F(1, 4) = 1.06 | 0.361 |
| | | | | CORT × KET | F(1, 4) = 0.21 | 0.669 |

**Supplementary Table S1** (continued)

| Panel | Culture | Parameter | Analysis | Effect / comparison | Statistic | p |
|---|---|---|---|---|---|---|
| 6f | Cortex | Events per active ROI | Two-way RM ANOVA | KET | $F(1, 4) = 4.79$ | 0.094 |
| | | | | CORT | $F(1, 4) = 0.87$ | 0.404 |
| | | | | CORT × KET | $F(1, 4) = 1.01$ | 0.372 |
| 6g | Cortex | Inter-event interval | Two-way RM ANOVA | KET | $F(1, 4) = 6.05$ | 0.070 |
| | | | | CORT | $F(1, 4) = 0.64$ | 0.469 |
| | | | | CORT × KET | $F(1, 4) = 0.32$ | 0.601 |
| 6h | Cortex | Amplitude of $Ca^{2+}$ transients | Two-way RM ANOVA | CORT | $F(1, 4) = 6.16$ | 0.068 |
| | | | | KET | $F(1, 4) = 9.60$ | 0.036 |
| | | | | CORT × KET | $F(1, 4) = 6.49$ | 0.063 |
| 6i | Cortex | Global synchronization index | Two-way RM ANOVA | KET | $F(1, 4) = 18.88$ | 0.012 |
| | | | | CORT | $F(1, 4) = 0.83$ | 0.413 |
| | | | | CORT × KET | $F(1, 4) = 0.26$ | 0.640 |
| Fig. S2a | Cortex | Assemblies per cell | Two-way RM ANOVA | CORT | $F(1, 4) = 18.32$ | 0.013 |
| | | | | KET | $F(1, 4) = 2.33$ | 0.202 |
| | | | | CORT × KET | $F(1, 4) = 0.001$ | 0.980 |
| Fig. S2b | Cortex | Participation index | Two-way RM ANOVA | KET | $F(1, 4) = 14.66$ | 0.019 |
| | | | | CORT | $F(1, 4) = 0.87$ | 0.404 |
| | | | | CORT × KET | $F(1, 4) = 0.27$ | 0.630 |
| Fig. S2c | Cortex | CV of transient amplitude | Two-way RM ANOVA | KET | $F(1, 4) = 21.80$ | 0.010 |
| | | | | CORT | $F(1, 4) = 0.001$ | 0.973 |
| | | | | CORT × KET | $F(1, 4) = 0.45$ | 0.540 |
| Fig. S2d | Cortex | Rise time | Two-way RM ANOVA | CORT | $F(1, 4) < 0.001$ | 0.996 |
| | | | | KET | $F(1, 4) = 0.27$ | 0.628 |
| | | | | CORT × KET | $F(1, 4) = 0.12$ | 0.749 |
| Fig. S2e | Cortex | Fall time | Two-way RM ANOVA | CORT | $F(1, 4) = 0.93$ | 0.389 |
| | | | | KET | $F(1, 4) = 0.61$ | 0.480 |
| | | | | CORT × KET | $F(1, 4) = 7.21$ | 0.055 |
| 7c | Hippocampus | ROIs per field | Two-way RM ANOVA | CORT | $F(1, 4) = 2.57$ | 0.184 |
| | | | | KET | $F(1, 4) = 2.37$ | 0.198 |
| | | | | CORT × KET | $F(1, 4) = 0.18$ | 0.692 |
| 7d | Hippocampus | Active ROIs (%) | Two-way RM ANOVA | KET | $F(1, 4) = 13.84$ | 0.020 |
| | | | | CORT | $F(1, 4) = 1.32$ | 0.315 |
| | | | | CORT × KET | $F(1, 4) = 1.59$ | 0.275 |

**Supplementary Table S1** (continued)

| Panel | Culture | Parameter | Analysis | Effect / comparison | Statistic | p |
|---|---|---|---|---|---|---|
| 7e | Hippocampus | Events per active ROI | Two-way RM ANOVA | CORT × KET | F(1, 4) = 10.05 | 0.034 |
| | | | | CORT | F(1, 4) = 2.40 | 0.196 |
| | | | | KET | F(1, 4) = 9.75 | 0.035 |
| | | | | Planned t-test: DMSO vs CORT | t(4) = 1.89 | 0.131 |
| | | | | Planned t-test: CORT vs CORT + KET | t(4) = 4.05 | 0.015 |
| | | | | Exploratory t-test: DMSO vs CORT + KET | t(4) = 0.42 | 0.694 |
| 7f | Hippocampus | Inter-event interval | Two-way RM ANOVA | CORT | F(1, 4) = 8.41 | 0.044 |
| | | | | KET | F(1, 4) = 4.25 | 0.108 |
| | | | | CORT × KET | F(1, 4) = 0.28 | 0.627 |
| 7g | Hippocampus | Amplitude of $Ca^{2+}$ transients | Two-way RM ANOVA | KET | F(1, 4) = 5.32 | 0.082 |
| | | | | CORT | F(1, 4) = 0.006 | 0.944 |
| | | | | CORT × KET | F(1, 4) = 0.18 | 0.695 |
| 7h | Hippocampus | Global synchronization index | Two-way RM ANOVA | KET | F(1, 4) = 19.65 | 0.011 |
| | | | | CORT | F(1, 4) = 0.32 | 0.600 |
| | | | | CORT × KET | F(1, 4) = 1.03 | 0.367 |
| Fig. S2f | Hippocampus | Assemblies per cell | Two-way RM ANOVA | KET | F(1, 4) = 7.69 | 0.050 |
| | | | | CORT | F(1, 4) = 0.39 | 0.566 |
| | | | | CORT × KET | F(1, 4) = 0.89 | 0.399 |
| Fig. S2g | Hippocampus | Participation index | Two-way RM ANOVA | KET | F(1, 4) = 19.13 | 0.012 |
| | | | | CORT | F(1, 4) = 0.04 | 0.857 |
| | | | | CORT × KET | F(1, 4) = 1.09 | 0.356 |
| Fig. S2h | Hippocampus | CV of transient amplitude | Two-way RM ANOVA | CORT | F(1, 4) = 1.62 | 0.273 |
| | | | | KET | F(1, 4) = 3.27 | 0.145 |
| | | | | CORT × KET | F(1, 4) = 0.40 | 0.563 |
| Fig. S2i | Hippocampus | Rise time | Two-way RM ANOVA | CORT | F(1, 4) = 30.48 | 0.005 |
| | | | | KET | F(1, 4) = 15.97 | 0.016 |
| | | | | CORT × KET | F(1, 4) = 15.24 | 0.017 |
| | | | | Planned t-test: DMSO vs CORT | t(4) = 0.86 | 0.436 |
| | | | | Planned t-test: CORT vs CORT + KET | t(4) = 1.72 | 0.160 |
| Fig. S2j | Hippocampus | Fall time | Two-way RM ANOVA | CORT × KET | F(1, 4) = 9.05 | 0.040 |
| | | | | CORT | F(1, 4) = 1.72 | 0.260 |
| | | | | KET | F(1, 4) = 1.57 | 0.279 |
| | | | | Planned t-test: DMSO vs CORT | t(4) = 0.50 | 0.642 |
| | | | | Planned t-test: CORT vs CORT + KET | t(4) = 0.92 | 0.410 |
